# “Convergent evolutionary selection unravels the genetic basis of audition in moths”

**DOI:** 10.64898/2026.07.08.736348

**Authors:** Scott D. Cinel, Quentin Flattmann, Chandra Earl, Emily Ellis, Jesse Barber, Natasha Mhatre, Yash Sondhi, Akito Y. Kawahara

**Author notes:** contributed equally (Scott D. Cinel, Quentin Flattmann, Yash Sondhi, Akito Y. Kawahara). **Corresponding authors**: Dr. Akito Kawahara, Dr. Yash Sondhi, Scott Cinel.

## Abstract

Hearing in Lepidoptera mediates a range of ecologically important behaviours, including mate communication, predator avoidance, and acoustic signalling. In moths, the evolution of predator-prey interactions with bats has further shaped hearing through a sensory arms race, with repeated co-option of auditory organs to detect and evade echolocating predators. Despite significant prior characterization of the neurophysiology and behaviour of hearing in moths, the genetic basis of hearing is poorly understood in most insects. In this study, we identify a core set of putative auditory genes in Lepidoptera using a combination of homology-based searches from *Drosophila* and evolutionary rate analyses. We find 56 genes present across all species and investigate whether gene copy number varies among non-hearing and hearing lineages and among 3 different ear types. We discovered seven genes associated with ear type and one with ear presence, but did not find significant losses in gene copy number in non-hearing species. We identified three genes (*btv*, *Dnai2*, and *nompB*) with strong evidence of selection in hearing clades and five genes with weaker evidence of selection. We discuss the potential roles of *btv, nompB,* and *Dnai2* in ciliary transport and the aging of hair cells, as well as the possibility of actively amplified hearing. Our study serves as a primer and resource for further gene mining and functional testing of auditory genes in moths and other insects.

## Introduction

Hearing is a vital sensory function found throughout insect species, often essential to successful mating, predator avoidance, and communication. It is thought to have evolved in 20 independent instances, with most insects (e.g. moths, crickets) possessing a dedicated tympanum [1], and with antennal ears being less common (e.g. flies, mosquitos). While such hearing organs are diverse in morphology, they are widely considered to have arisen from the same pre-existing proprioceptive or low-frequency vibration-sensitive mechanosensory chordotonal sensillae [1–4]. The ancestral state is thought to be a proprioceptive organ at the wing hinge which senses wing beats, and is still present in several moth lineages [5]. Among the simplest ears are the broadband pressure detecting ears found in moths and butterflies, lacewings, beetles, and mantids [6]. These ears function to detect ultrasonic echolocation signals of insectivorous bats and allow the animals to perform evasive flight or deploy other anti-predatory strategies in response to bat calls [7–12].

Lepidoptera, the second largest insect order, contains over 180,000 species of moth and butterfly [13]. Multiple separate evolutions of tympanal organs have occurred in Lepidoptera with varying anatomical placement, including on the thorax, abdomen, wings, and mouthparts [14]. Prior work in moth hearing has suggested that the metathoracic tympanal ear arose once in the phylogeny and is the now extinct ancestor of the notodontid and noctuid moths [6, 14, 15]. The genes underlying mechanosensation in chordotonal organs likely provided the basis for a gradual evolutionary transition towards tympanal hearing as opposed to it being a de novo innovation [16].

Comparisons between the hearing systems of *Drosophila* vinegar flies versus Lepidoptera reveal both similarities and differences in anatomical structure, suggesting the presence of shared genetic components underlying auditory function [17–19]. In *D. melanogaster,* many genes with roles in the auditory function of ciliated mechanosensory neurons have been described including those encoding mechanotransduction channels (e.g. *TRP*, *Piezo*), ion channels and GPCR regulators, ciliary structure (e.g. dynein, kinesin, *RFX*), cytoskeletal and mechanical coupling (e.g. actin, ankyrin), and developmental patterning (e.g. *atonal*, *salm*, *engrailed*) [18, 20–22]. Targeted knockout and mutation of these genes in *Drosophila* result in defects in auditory organ development and/or function [21].

One approach to understanding the evolution of hearing among Lepidoptera is to examine the evolution of gene families and the rates of gene evolution associated with a given hearing organ phenotype [23]. This approach has been fruitful in other sensory systems such as vision, olfaction, and wing patterning [24–28].

Lepidoptera includes species with diverse tympanal organs, likely derived through separate evolutionary origins. Hearing and sound production in moths have been subject to extensive study, particularly with regard to bat-moth interactions [29–33], morphology, neurophysiology, and behavior. Despite the diverse implementations and functions of hearing in Lepidoptera, the order’s genetic basis of hearing remains largely unexplored. The few Lepidoptera-specific sensory-gene surveys [35–36] that have been conducted generally lack broad comparative genomic sampling, hearing versus non-hearing contrasts, or tests of selection and copy-number evolution. In contrast, numerous genes involved in hearing have been described in the model organism Drosophila melanogaster [21]. These genes represent a valuable and practical candidate set for initial investigations into auditory evolution in other insects. However, the relevance of these genes in Lepidoptera remains unclear given the substantial differences in morphology and anatomy between the two orders, and the fact that the lineages are estimated to have diverged roughly 284 Ma [34].

We endeavour to bridge this gap in the understanding of moth hearing by focusing on gene homology and evolutionary rate analyses, aiming to identify a set of candidate genes that may underlie hearing in Lepidoptera. Given that lepidopteran tympanal ears are thought to have evolved from pre-existing vibration-sensitive chordotonal organs, we hypothesize that hearing evolution in Lepidoptera involved primarily the modification of conserved mechanosensory genes rather than the use of an entirely novel set of hearing genes. Under this hypothesis, we predict that *Drosophila* hearing-related genes will be broadly conserved across both hearing and non-hearing lepidopteran lineages, but that hearing lineages will show distinct signatures of copy-number variation or altered selective pressure in a subset of these conserved genes. Using a dataset of 19 high-quality lepidopteran genome assemblies, representing hearing lineages with multiple different ear morphologies as well as non-hearing lineages, we focus on identifying homologs of *Drosophila* hearing-related genes and assessing their evolutionary dynamics. The hearing taxa span in this dataset include three tympanal ear morphologies: pyraloid abdominal ears, noctuoid metathoracic ears, and the atypical prothoracic ear of *Operophtera brumata* (Supplementary Table 3). First, we aim to determine whether all *Drosophila* hearing-related genes are present in these 19 Lepidoptera or if some have been lost in non-hearing lepidopteran clades. Next, we seek to investigate whether hearing genes exhibit differing patterns of positive selection between clades. Finally, we aim to characterize the functionality of the sites under selection of the hearing genes we identify by mapping these sites to PFAM domains. Thus, through this study, we seek to begin identifying the gene repertoire involved in hearing in Lepidoptera, providing valuable insights into the evolutionary history, molecular pathways, and functional adaptations that underlie the diversity of this sensory modality among butterflies and moths.

## Results

### Hearing gene copy number is more strongly associated with ear type than ear presence

A comparison across 3 different ear types and ear presence versus absence was performed to identify patterns in gene copy number variation of orthologous hearing genes among the taxa studied. The hearing taxa in this study represent three lepidopteran tympanal ear types: pyraloid abdominal ears, noctuoid metathoracic ears, and the atypical prothoracic ear of *Operophtera brumata*.

#### (i) Hearing vs Non-Hearing Copy Number Variation (CNV)

Increased copy number in the *bmcp* gene was found to be significantly associated with ear presence with a p-value of 0.019, as indicated by the Mann-Whitney test (Supplementary Table 4). Notably, the remaining 55 genes showed no statistically significant variation in copy number between hearing and non-hearing species. *bmcp* is an unusual hearing-associated gene in *D. melanogaster*, given that it is involved in mitochondrial metabolism. However, *bmcp* knockout results in moderate auditory impairment, suggesting *bmcp* may play a role in auditory mechanoreceptor cell metabolism in insects. [21].

#### (ii) Hearing Organ Type CNV

Eight genes were found to vary significantly in copy number by ear type via a Chi-Square test (Supplementary Table 5). For these eight genes, we performed pairwise Mann-Whitney tests to compare copy number between each pair of ear-types to assess for specific differences (Supplementary Table 6). However, all significant pairwise comparisons included the type 3 ear type, the atypical prothoracic tympanal ear, and are ultimately limited in their implications given that this group contains only one species.

### Evidence for altered selection pressure in hearing vs. non hearing species

Given that orthologs of the genes selected in this study were present in the majority of species, an analysis of selection rate differences between hearing and non-hearing species was performed using 3 different selection tests: aBSREL, BUSTED, and Contrast-FEL. We found different genes under selection among each test, with 34/56 genes under selection according to at least 1 test (Supplementary Table 7). We found 9 genes with evidence of selection corroborated by at least two tests, and of these, *btv*, *nompB*, and *Dnai2* showed evidence of selection in all 3 tests.

### Rates of evolution and sensitivity to stringency tests

#### (i) BUSTED Selection Test

The Likelihood Ratio Test (LRT) is an output metric of BUSTED which computes the probability of episodic diversifying selection and revealed 10 genes with a high probability of this condition with p-value < 0.05, with an additional 8 genes indicated with a p-value < 0.10 (Figure 3A).

**Figure 2:**
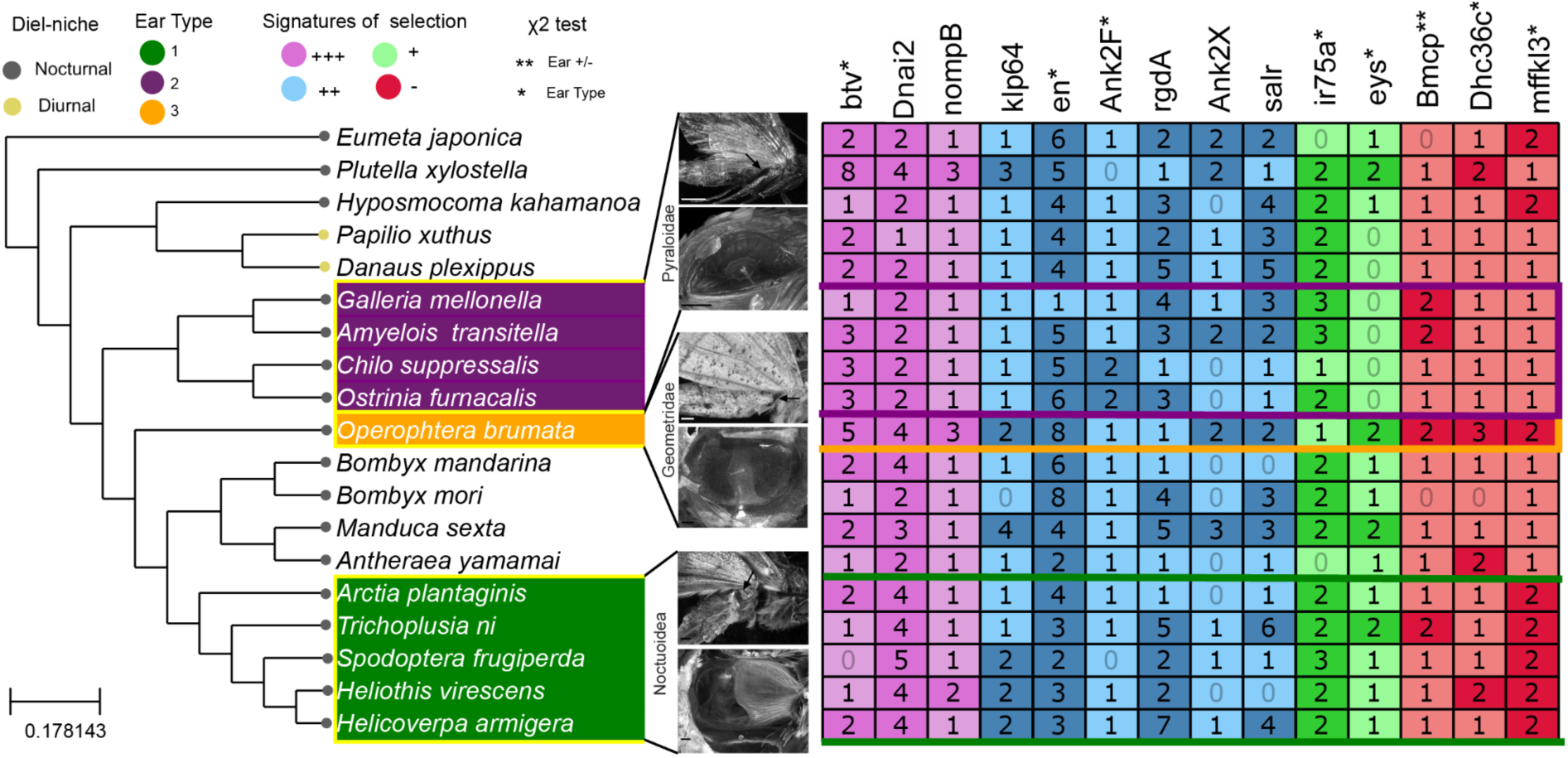
Number of gene copies identified for orthologous hearing genes derived from *Drosophila melanogaster* across hearing and non-hearing lepidopteran clades. For illustrative purposes a cladogram has been shown, but trees with branch lengths were used for all analyses.

**Figure 2:**
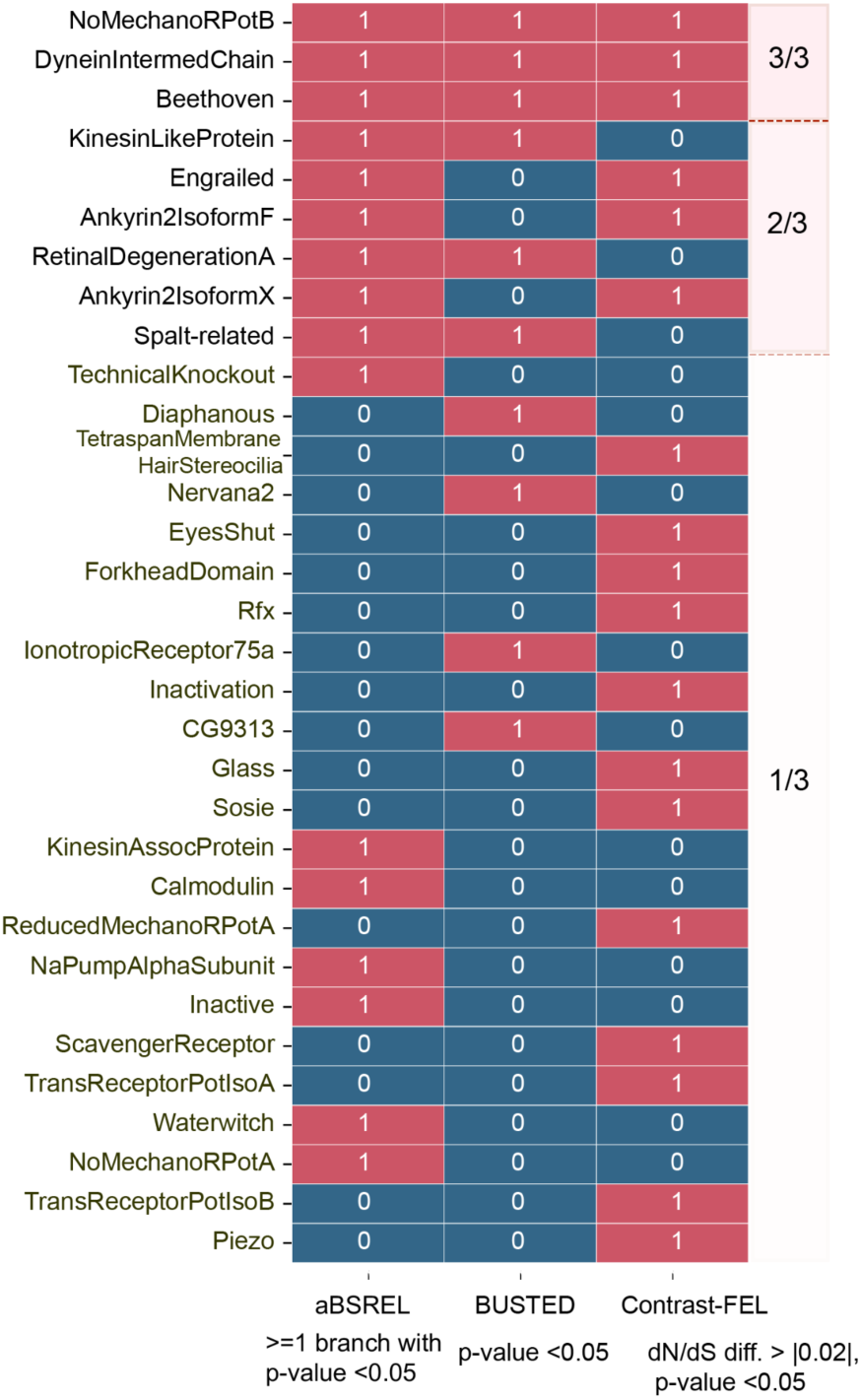
Genes under selection in hearing relative to non-hearing lepidopteran clades using different selection tests (aBSREL, BUSTED, Contrast-FEL; red indicates evidence of selection, blue indicates no evidence of selection).

#### (ii) Contrast-FEL Selection Test

In cases where a mean ω for hearing and non-hearing branches had been calculated, we used Contrast-FEL testing to identify which genes were under positive or negative selection with respect to each lineage. This follows prior comparative applications of Contrast-FEL in which the direction and number of codon sites with elevated dN/dS are used to infer lineage- or phenotype-associated shifts in selective pressure [37–38]. Notably, *Dnai2* was under positive selection along with Anykrin (*ANK*) and inactive (*iav*), but all other genes were under negative selection. In this case, the background was defined as non-hearing species and the calculation consisted of the dN/dS of hearing species – dN/dS of non-hearing species.

Contrast-FEL analyses identified 35 of 56 genes with significant codon sites differing in selective pressure between hearing and non-hearing lineages. Of these, 27 genes showed a greater number of sites with elevated ω in non-hearing taxa, whereas 8 genes showed more sites with elevated ω in hearing taxa (Supplementary Tables 1-2). This suggests that although hearing-associated genes are broadly conserved across Lepidoptera, lineage-specific shifts in constraint and adaptive evolution are widespread among non-hearing species. To further characterize selective differences identified by Contrast-FEL, we examined whether codon sites with significantly different selection rates among hearing and non-hearing lineages were disproportionately enriched for elevated ω in either lineage. Ankyrin 2 isoform X (*ANK2X*) exhibited a strong excess of sites with elevated ω in hearing taxa (104 sites vs. 26), suggesting lineage-specific adaptive refinement or reduced constraint associated with auditory specialization. In contrast, *engrailed* showed more sites with elevated ω in non-hearing taxa (17 sites vs. 9), consistent with divergent selective pressures in non-hearing lineages. Other notable genes, *btv* and *cut*, displayed numerous significant codon sites but lacked strong directional differences between hearing and non-hearing clades.

#### (iii) aBSREL Selection Test

Given that aBSREL tests for episodic diversifying selection on individual branches, we used it to ask whether hearing-associated lineages showed branch-specific evidence of positive selection and whether those branches corresponded to ear type. Of the 9 non-hearing species, aBSREL suggested no positive selection of any of the selected genes with p-value < 0.05. Notably, we found 7 genes with evidence of episodic selection in *O. brumata* while *S. frugiperda* had 4. *O. brumata* is the only species in this study with atypical tympanal ears present on the prothorax, near the base of the forelegs (Supplementary Table 3). These results suggest that the atypical tympanal ear structure drew the most on existing ancestral genes. Genes appearing with episodic selection in more than one species do not appear to be correlated to ear type. For example, Ankyrin 2 isoform F (*ANK2F*) appears to have evolved under episodic positive selection in *G. mellonella*, *T. nii*, and *O. brumata*, each of which displays an anatomically distinct ear type.

**Figure 3:**
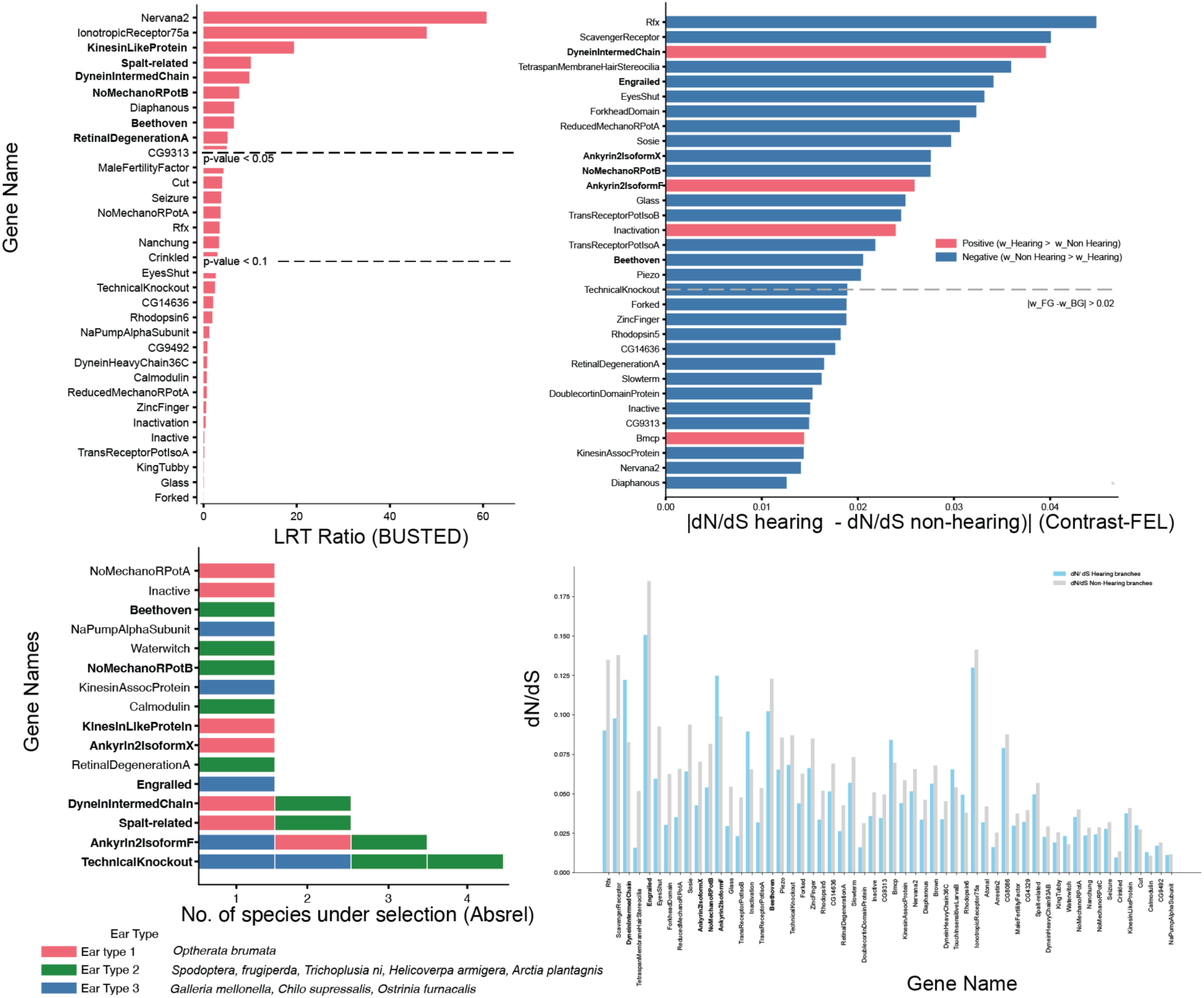
Genes under selection for A: BUSTED, B: Contrast-FEL, C: aBSREL and D: estimated dN/dS values for hearing and non-hearing branches.

### Selection on Conserved PFAM Domains

No clear pattern of increased selection on any predicted PFAM domains in the genes *ANK2F*, *ANK2X*, *btv*, *cut*, and *engrailed* was found. This may indicate conservation of critical residues or adaptive changes. However, the resolution of PFAM domain predictions may be too coarse to clearly detect domain-specific trends.

## Methods

### Identification of Drosophila hearing gene set using previous literature

We first combed the *Drosophila* literature to identify genes related to hearing. Prior research was drawn on to produce a list of 56 genes, 52 of which have Gene Ontology annotations pertaining to the transduction of sound specifically (‘sensory perception of sound’ - (GO:0007605) [21–22].

**Figure 4.**
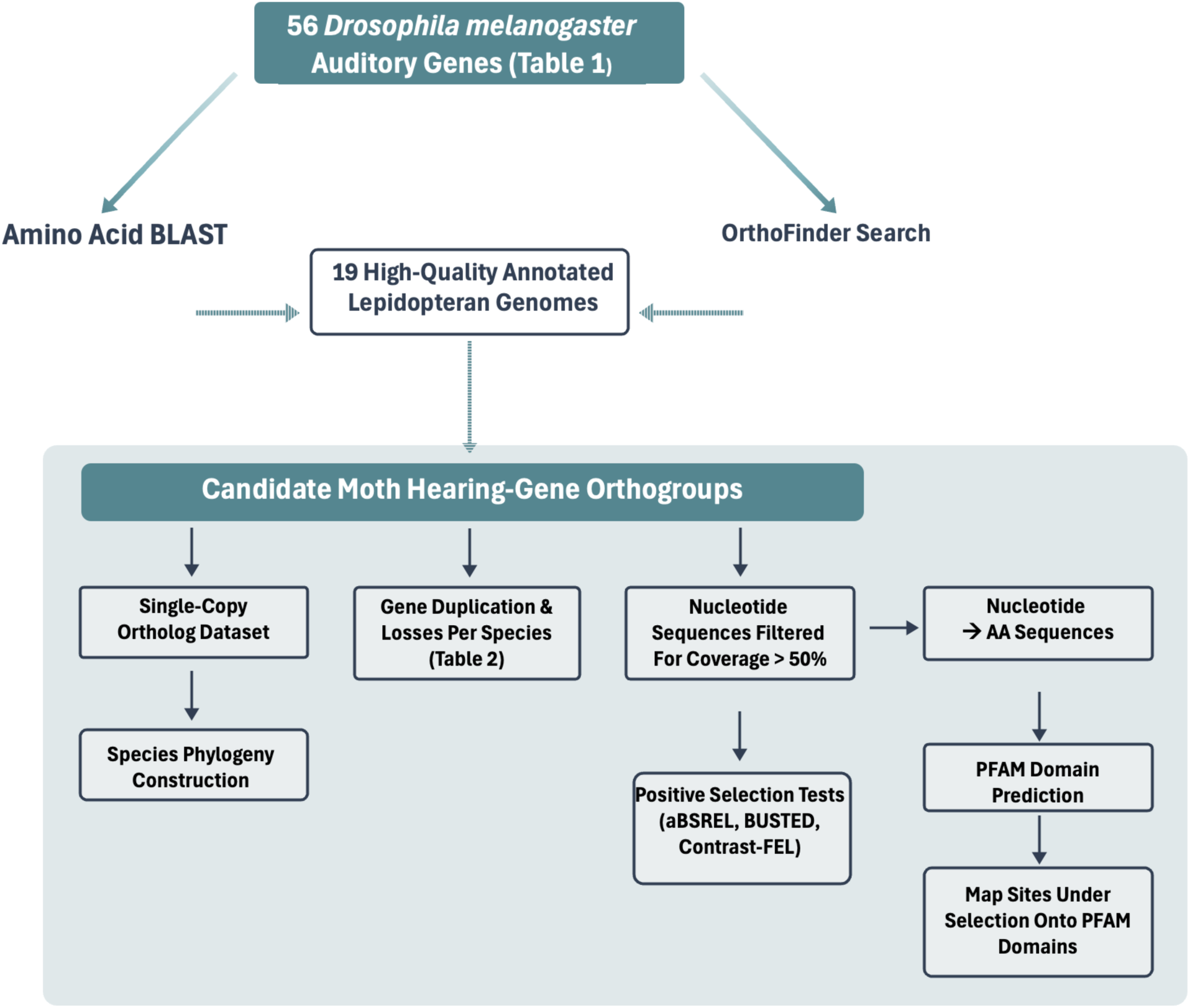
Bioinformatic workflow for identifying candidate hearing genes in Lepidoptera and testing evolutionary patterns of gene loss, duplication, and selection. Starting from 56 *Drosophila melanogaster* auditory genes (Table 1), orthologous sequences were identified in 19 high-quality annotated lepidopteran genomes using amino-acid BLAST and OrthoFinder. Candidate orthogroups were then used for (i) copy-number and presence/absence analysis (Table 2), (ii) species phylogeny construction from single-copy orthologs, (iii) positive selection tests (aBSREL, BUSTED, Contrast-FEL), and (iv) mapping of sites under selection onto predicted PFAM domains.

### Moth and Drosophila orthologous protein dataset generation with OrthoFinder

We identified 19 species of Lepidoptera with published, high-quality, annotated genomes, among which 3 different hearing organ types have evolved [39–40] (Supplementary Table 3).

Using the translated *Drosophila melanogaster* reference sequence for each of the 56 hearing genes as the query, we then used OrthoFinder v.2.4.0 with default parameters [41] to designate possible orthologous sequences from each of the lepidopteran genomes. Predicted proteomes from 19 lepidopteran species and a curated *D. melanogaster* hearing-gene bait set were analyzed jointly to identify orthogroups containing candidate hearing-related genes. A total of 65 bait-associated orthogroups were recovered, after which duplicate or sparsely represented groups were excluded prior to downstream analyses.

### Sequence Alignment & Gene Tree Construction

We then used each set of orthologous lepidopteran sequences pertaining to a particular gene, along with the *Drosophila* reference sequence, to align using MAFFT v.7.245 [42] and proceeded to construct gene trees using IQTree v.2.4.0 [43]. At this stage, we pruned tree tips from ingroups with very long branch lengths, realigned sequences, and rebuilt trees. The topology of each resulting tree was then manually verified to ensure that the orthologous sequences identified by OrthoFinder were in fact representative of the same gene and not errant matches. Specifically, we selected for further analysis only those sequences that were within the same clade or in clades sister to the reference *Drosophila* sequence. After this initial filtration, we retrieved the corresponding nucleotide coding sequences from the original species-specific coding sequence (CDS) files for each retained amino acid sequence.

We also removed any sequence that whose length was less than 50% of that of the consensus sequence in the resulting alignment, as identified in Geneious Prime v.2022.1.1 [44], and in cases where OrthoFinder provided multiple sequences from a given species for a specific gene, we selected the sequence best matching the consensus because selection tests are often limited to testing a single gene sequence per species (Supplementary Methods 1). Sequences were translated via the invertebrate translation table in Geneious and trimmed to the position before their first stop codon. Sequences with more than five stop codons were removed.

### Testing for selection on hearing genes

We then used HyPhy v.2.5.8 to implement tests of gene-wide (Branch-site Unrestricted Statistical Test for Episodic Diversification [BUSTED]), lineage-specific (adaptive Branch-site Random Effects Likelihood test for episodic diversification [aBSREL]), and site-specific (Contrast-Fixed Effects Likelihood; Contrast-FEL) signatures of positive selection. For Contrast-FEL, these signatures are determined according to dN/dS estimates, or the ratio of non-synonymous to synonymous amino acid substitutions present in each sequence [45]. We considered only those genes found to have evolved under positive selection by all three statistical tests as meaningful results.

We interpreted aBSREL results as evidence of episodic diversifying selection on individual branches, rather than evidence for uniform positive selection across the entire gene tree. This process follows prior interpretation methods such as the branch-site framework described by Smith et al. [46]. Similar foreground-branch interpretations of aBSREL have been used in comparative genomic studies testing clade or phenotype-associated selection, including analyses of high-elevation and direct-developing frog lineages [47].

Contrast-FEL was interpreted as a comparative site-wise test of differential selective pressure between predefined branch classes, following prior applications that compare dN/dS across focal and background lineages or phenotype-defined taxa [37, 38].

### Mapping of Significant Sites Identified By Contrast-FEL To PFAM Domains

To further characterize the functionality of sites found to be under selection, we mapped significant sites as identified by Contrast-FEL with p-value < 0.05 to predicted PFAM domains in *Bombyx mori* for five genes: *ANK2F*, *ANK2X*, *btv*, *cut*, and *engrailed*. PFAM domains were predicted against HMMER searches of the Pfam-A library. A filter was applied to keep only PFAM hits of high-confidence (E-value < 1e^-6^), and significant Contrast-FEL aligned coordinate sites were mapped to unaligned *B. mori* target gene sequences. Within each PFAM domain, we counted significant sites showing elevated selection, interpreted as evidence consistent with adaptive evolution, separately from sites showing reduced selection, interpreted as evidence of purifying selection. Their relative enrichment was calculated as the ratio between the percent of sites marked as significant in the domain compared to the percent in the overall gene.

## Discussion

We identified a conserved core set of 56 candidate hearing-related genes across Lepidoptera and found that they were broadly conserved in non-hearing species. Given the independent evolution of multiple lepidopteran ear types, we expected lineages with similar auditory phenotypes to show convergent patterns of gene gain or loss, analogous to sensory gene-family turnover observed in other systems, including opsins in vision and olfactory receptors in olfaction. However, most candidate genes were found to be retained across taxa with relatively modest copy-number differences. Several genes were linked to ear type with the most significant including *btv*, *dhct36c*, *engrailed*, *eys*, *ir75a*, *mffkl3*, *ANK2F*, and *bmcp*. Across multiple evolutionary models, a subset of genes—including *btv*, *nompB*, and *Dnai2*—showed robust signatures of positive selection in hearing lineages. The selection tests used in this study, aBSREL, BUSTED, and Contrast-FEL, revealed 10, 11, and 17 genes under selection, respectively. Ultimately, only a limited subset of genes showed significant associations with ear presence/absence and ear type, and these effects were not characterized by lineage-specific gene family turnover on a large scale.

This suite of genes has been generally conserved across species, likely due to their dual role in mechanosensation and hearing. Significantly, the three genes under robust selection are also directly implicated in ciliary structure and transport within mechanosensory neurons. These results suggest that the evolution of hearing in Lepidoptera is driven not by the emergence of novel genes, but by the modification of a conserved mechanosensory toolkit.

In *Drosophila*, mechanotransduction occurs in the ciliary dendrites of auditory neurons, hair-like sensory projections extending from auditory neurons in the Johnston’s organ. All three of the genes consistently identified to be under positive selection in this study, including *nompB*, *btv*, and *Dnai2*, are typically localised to the ciliary dendrite of auditory neurons in the Johnston’s organ. *btv* and *nompB* are associated with interflagellar transport (IFT), and also serve essential functions in the development of normal auditory chordotonal mechanosensory neurons, specifically of the ciliated dendrite [48–50]. *btv* encodes a motor protein, a cytoplasmic dynein, that moves in the retrograde direction along the ciliated dendrite [48–49]. In *btv* mutants, the ciliated dendrite takes on a malformed structure and mutant flies have impaired auditory function in early life which progressively worsens with age [48–49]. These findings are congruent with the idea that retrograde IFT is important for the removal of dysfunctional proteins and other molecules [48]. *nompB* is an intraflagellar transport protein associated with an anterograde motor. The dendritic cilia in the chordotonal organs of *nompB* mutants are almost completely absent and unable to produce potentials in response to sound [50].

*Dnai2* codes for the outer dynein arm expressed in the proximal part of the ciliary dendrite of *Drosophila* auditory neurons [51]. *Dnai2* mutants do not display any apparent developmental defects and develop a normal cilium, except for a missing dynein arm [51]. However, *Dnai2* mutants exhibit lowered neuronal sensitivity to sound and lack the antennal or neuronal non-linearity associated with active auditory amplification in *Drosophila* [51–52]. These prior findings establish that axonemal dynein-driven ciliary motility is required for active auditory amplification in *Drosophila* [51], while otoacoustic emissions from noctuoid moth tympanal organs and other simple insect ears suggest that active auditory mechanics may also occur in moth hearing [53–58].

This study is among the first to attempt to untangle the complicated auditory gene system underlying hearing in Lepidoptera and provides valuable insight into a set of putative hearing genes. However, confirmation of these findings will require functional testing, including RNA-sequencing, protein localisation, and genetic manipulation [59]. Our findings support a model in which insect hearing evolves through selection on pre-existing sensory machinery, providing a genetic framework for repeated, convergent origins of auditory systems.

## Acknowledgements

We thank members of the Kawahara lab, the high-performance cluster at the University of Florida, and the Ohio State Cluster, NSERC Discovery (Grant no. 687216), and an NSERC Canada research chair (Grant no. 693206). This research was conducted with Government support under and awarded by DoD, Air Force Office of Scientific Research, National Defense Science and Engineering Graduate (NDSEG) Fellowship, 32 CFR 168a.

## Supplementary Tables

**Supplementary Table 1.** These genes exhibit a greater number of codon sites with elevated nonsynonymous substitution rates in hearing lineages relative to non_hearing lineages (ω_hearing_ > ω_non-hearing_), as indicated by Contrast-FEL results. The enrichment value indicates the portion of sites with ω_hearing_ > ω_non-hearing_ over the total number of significant sites.

| Gene | Number of Sites:<br>$\omega_{\text{hearing}} > \omega_{\text{non-hearing}}$ | Number of Sites:<br>$\omega_{\text{hearing}} < \omega_{\text{non-hearing}}$ | Total Significant Sites<br>($p_{\text{Overall}} > 0.05$) | Enrichment |
| --- | --- | --- | --- | --- |
| NaPumpAlphaSubunit | 6 | 0 | 6 | 1 |
| DyneinIntermediate Chain | 33 | 2 | 35 | 0.942857143 |
| Rhodopsin6 | 15 | 1 | 16 | 0.9375 |
| DoublecortinDomainProtein | 11 | 1 | 12 | 0.916666667 |
| Forked | 21 | 5 | 26 | 0.807692308 |
| Ankyrin2IsoformX | 104 | 26 | 130 | 0.8 |
| TouchInsensitiveLarvaB | 13 | 6 | 19 | 0.684210526 |
| Glass | 6 | 3 | 9 | 0.666666667 |

**Supplementary Table 2.**
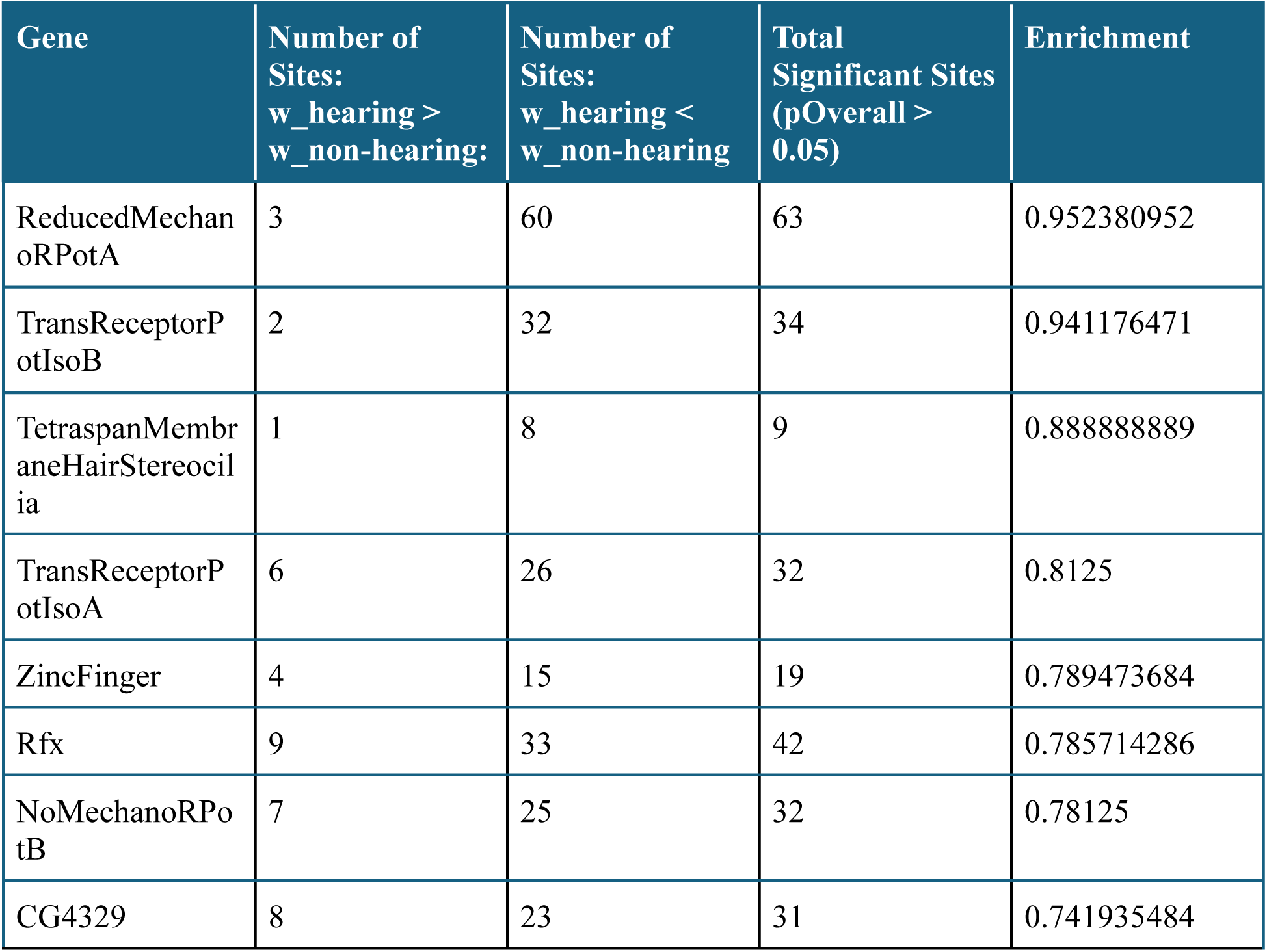
These genes exhibit a greater number of codon sites with elevated nonsynonymous substitution rates in non-hearing lineages relative to hearing lineages (ω_non-hearing > ω_hearing), as indicated by Contrast-FEL results. The enrichment value indicates the proportion of sites with ω_hearing < ω_non-hearing over the total number of significant sites.

| Gene | Number of Sites:<br>w_hearing > w_non-hearing: | Number of Sites:<br>w_hearing < w_non-hearing | Total Significant Sites<br>(pOverall > 0.05) | Enrichment |
| --- | --- | --- | --- | --- |
| ReducedMechanoRPotA | 3 | 60 | 63 | 0.952380952 |
| TransReceptorPotIsoB | 2 | 32 | 34 | 0.941176471 |
| TetraspanMembraneHairStereocilia | 1 | 8 | 9 | 0.888888889 |
| TransReceptorPotIsoA | 6 | 26 | 32 | 0.8125 |
| ZincFinger | 4 | 15 | 19 | 0.789473684 |
| Rfx | 9 | 33 | 42 | 0.785714286 |
| NoMechanoRPotB | 7 | 25 | 32 | 0.78125 |
| CG4329 | 8 | 23 | 31 | 0.741935484 |

**Supplementary Table 3.** This table shows the species designations of 19 Lepidoptera genomes used in this study. Ear types 0, 1, 2, and 3 indicate non-hearing taxa, pyramid abdominal tympanal ears, paired metathoracic tympanal ears, and prothoracic/foreleg tympanal ears, respectively.

| Genus | Species | Superfamily | Family | Subfamily | Tipnames | Ear Type | Butterfly/Moth |
| --- | --- | --- | --- | --- | --- | --- | --- |
| Eumeta | japonica | Tineoidea | Psychidae | Oiketicinae | Eumeta_japonica | 0 | M |
| Plutella | xylostella | Yponomeutoidea | Plutellidae | Plutellinae | Plutella_xylostella | 0 | M |
| Hyposmocoma | kahamanoa | Gelechioidea | Cosmopterigidae | Cosmopteriginae | Hyposmocoma_kahamanoa | 0 | M |
| Papilio | xuthus | Papilionoidea | Papilionidae | Papilioninae | Papilio_xuthus | 0 | B |
| Danaus | plexippus | Papilionoidea | Papilionidae | Danainae | Danaus_plexippus | 0 | B |
| Operophtera | brumata | Geometroidea | Geometridae | Larentiinae | Operophtera_brumata | 3 | M |
| Bombyx | mandarina | Bombycoidea | Bombycidae | Bombycinae | Bombyx_mandarina | 0 | M |
| Bombyx | mori | Bombycoidea | Bombycidae | Bombycinae | Bombyx_mori | 0 | M |
| Manduca | sexta | Bombycoidea | Saturniidae | Sphinginae | Manduca_sexta | 0 | M |
| Antheraea | yamamai | Bombycoidea | Saturniidae | Saturniinae | Antheraea_yamamai | 0 | M |
| Trichoplusia | ni | Noctuoidea | Noctuidae | Plusiinae | Trichoplusia_ni | 2 | M |
| Spodoptera | frugiperda | Noctuoidea | Noctuidae | Noctuinae | Spodoptera_frugiperda | 2 | M |
| Heliothis | virescens | Noctuoidea | Noctuidae | Heliothinae | Heliothis_virescens | 2 | M |
| Helicoverpa | armigera | Noctuoidea | Noctuidae | Heliothinae | Helicoverpa_armigera | 2 | M |
| Arctia | plantaginis | Noctuoidea | Erebidae | Arctiinae | Arctia_plantaginis | 2 | M |
| Galleria | mellonella | Noctuoidea | Noctuidae | Gallerinae | Galleria_mellonella | 1 | M |
| Amyelois | transitella | Pyraloidea | Pyralidae | Phycitinae | Amyelois_transitella | 1 | M |
| Chilo | suppressalis | Pyraloidea | Crambidae | Crambinae | Chilo_suppressalis | 1 | M |
| Ostrinia | furnacalis | Pyraloidea | Crambidae | Pyraustinae | Ostrinia_furnacalis | 1 | M |

**Supplementary Table 4.**
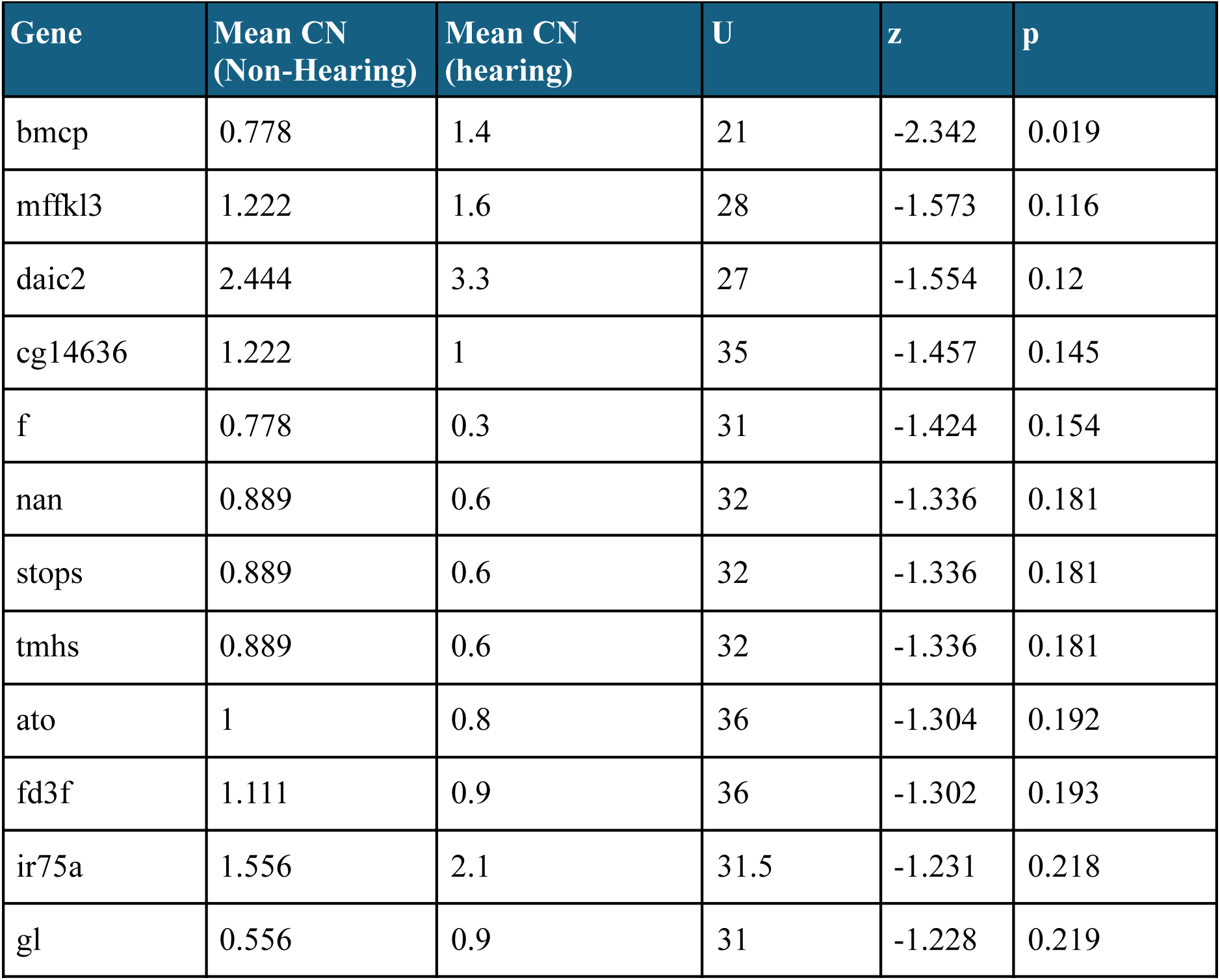
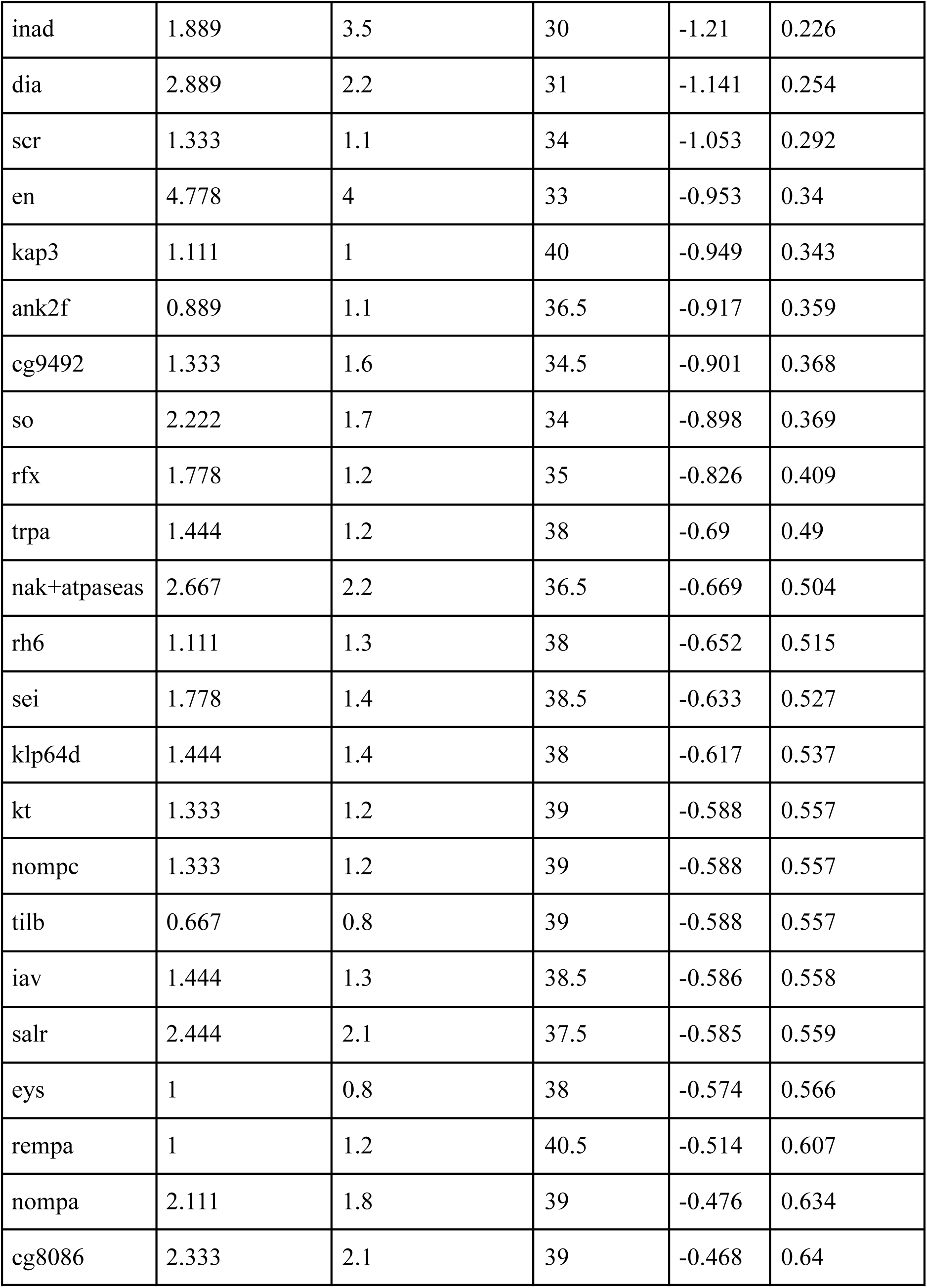
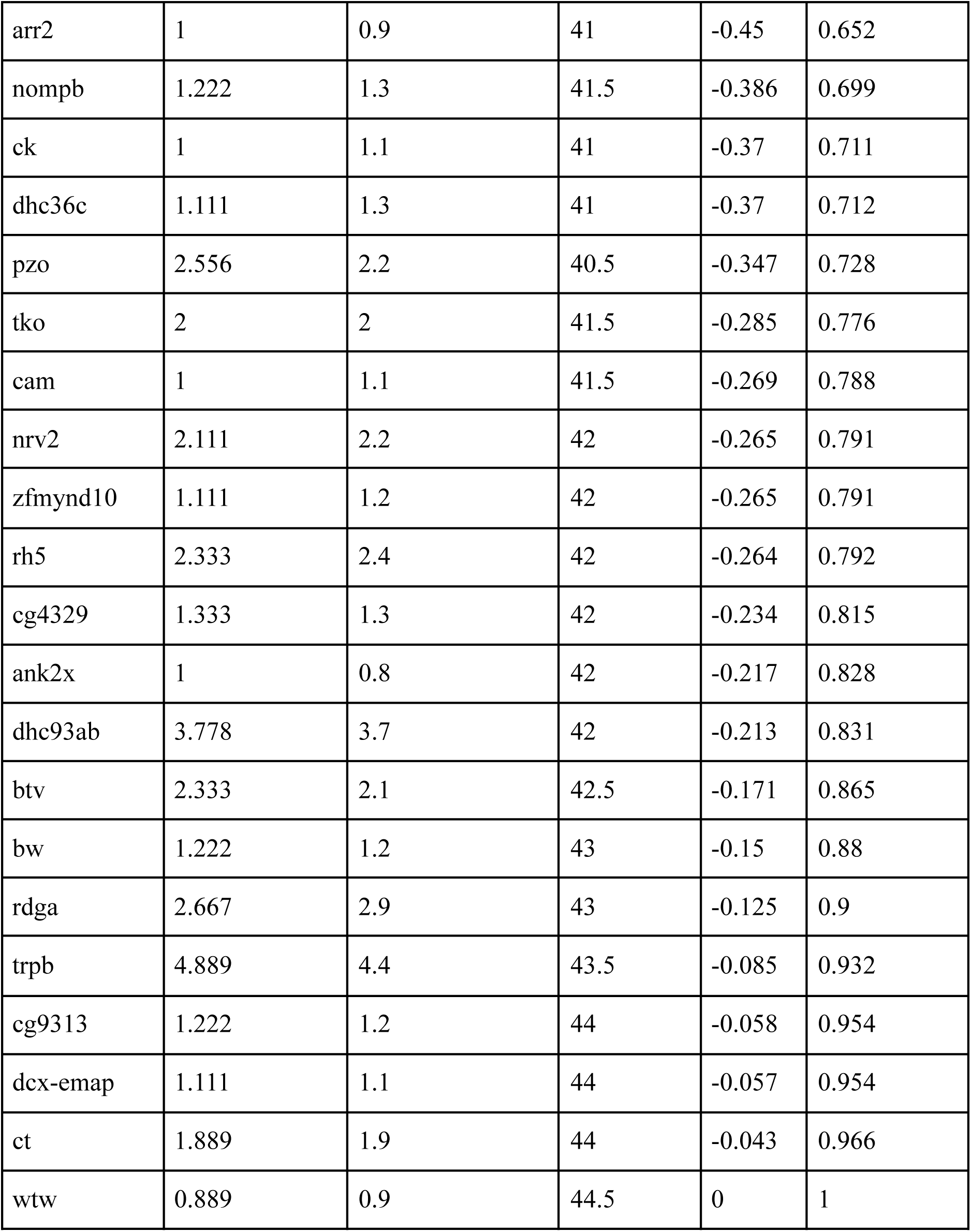
These results from Mann-Whitney U tests compare gene copy number (CN) between non-hearing and hearing taxa. For each gene, statistical significance outputs are shown which compare CN between groups (U statistic, standardized z-score, two-sided p-value).

**Supplementary Table 5.**
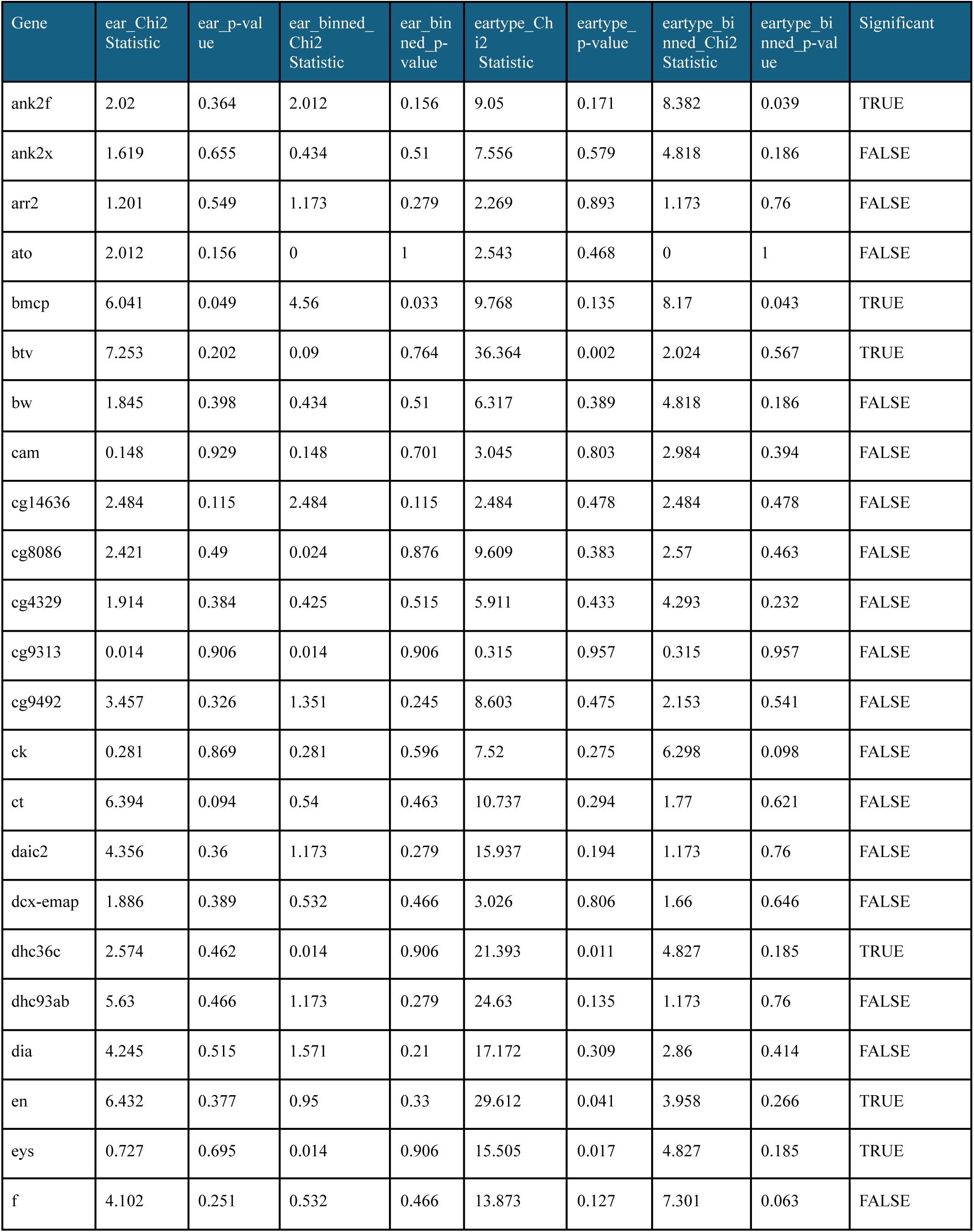
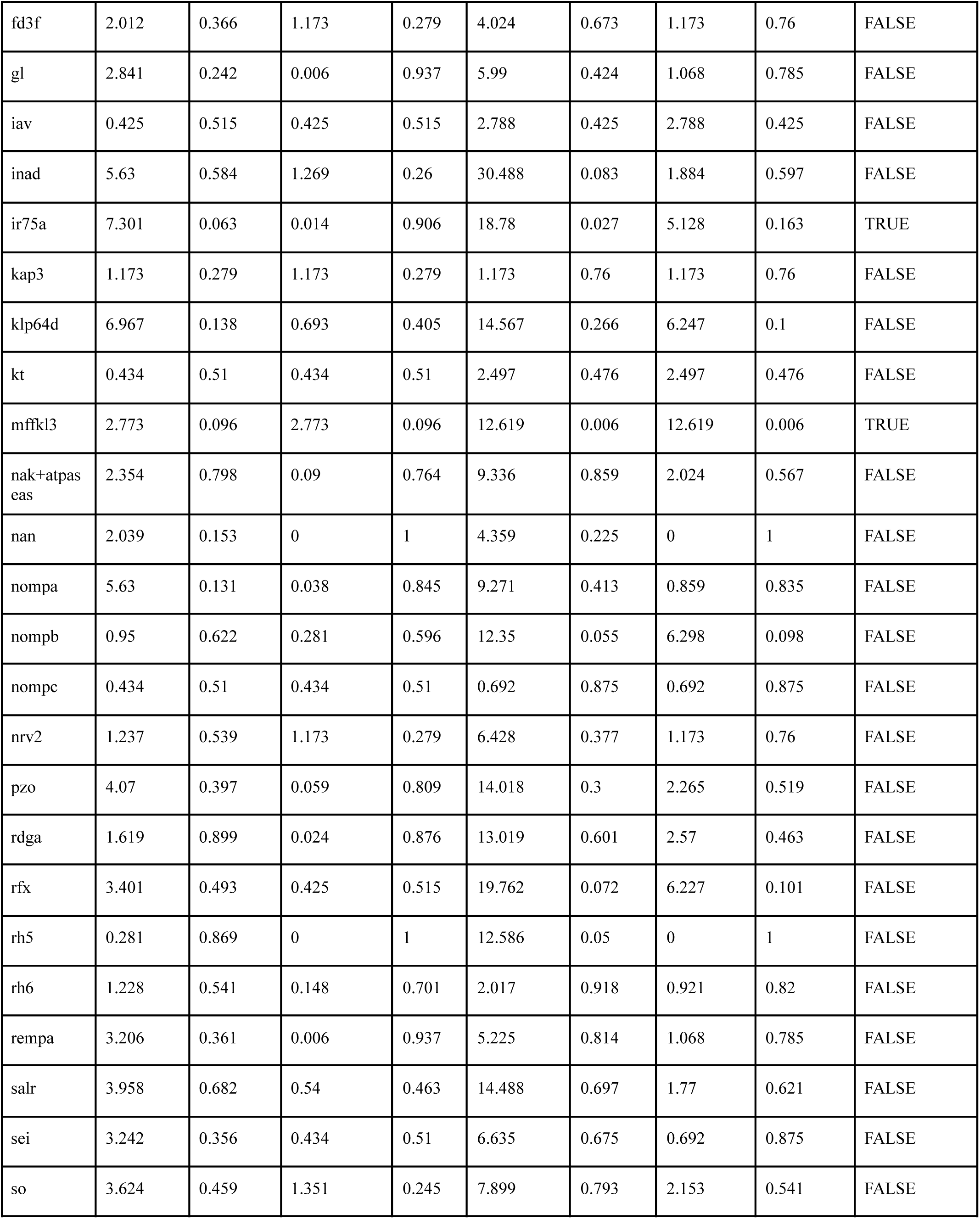
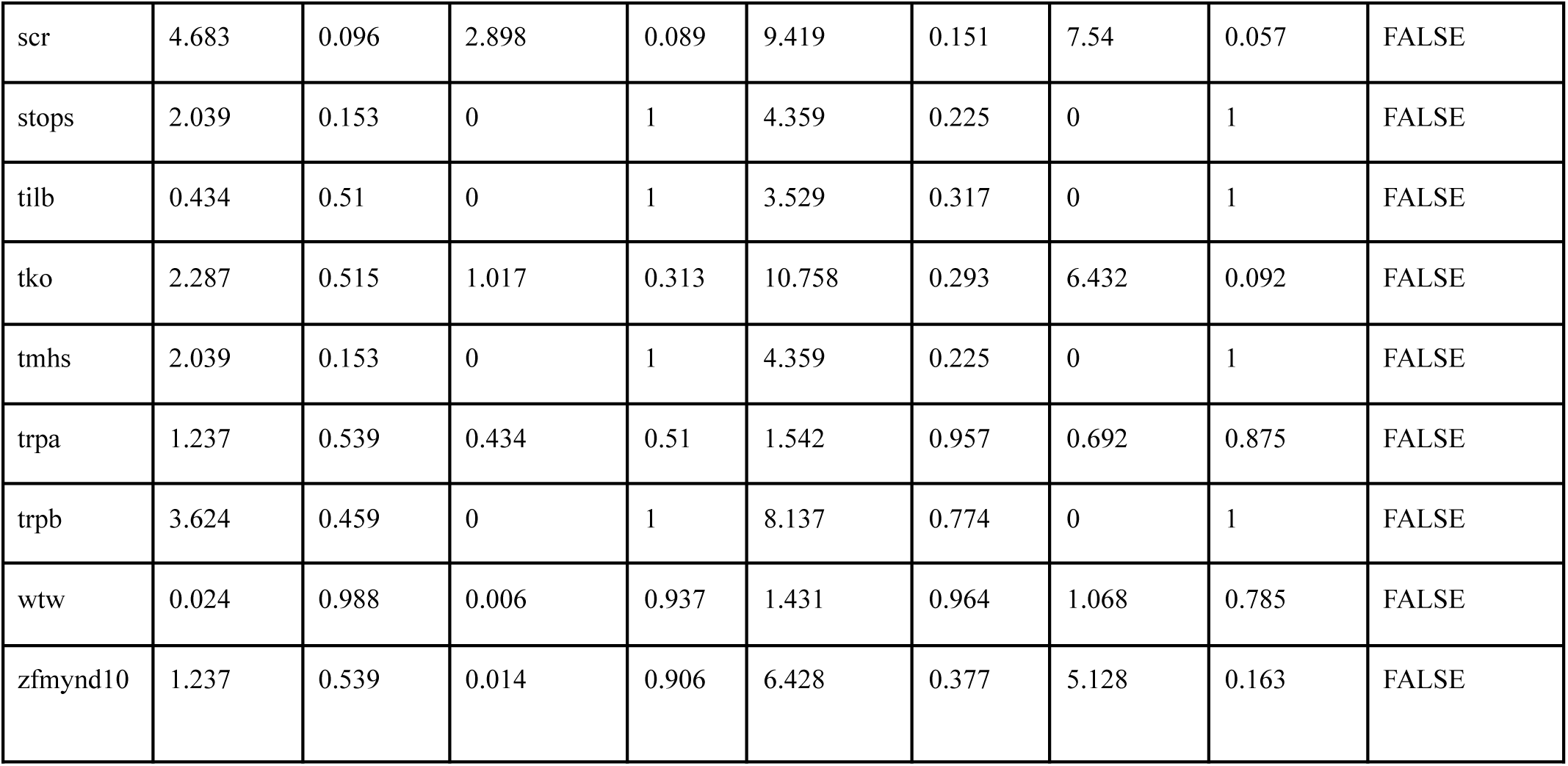
These results from chi-square associated tests evaluate the significance of copy number (CN) differences between hearing phenotypes across the 19 species of Lepidoptera. Ear tests are a comparison of non-hearing and hearing taxa whereas ear type tests compare the three hearing organ morphologies.

**Supplementary Table 6.**
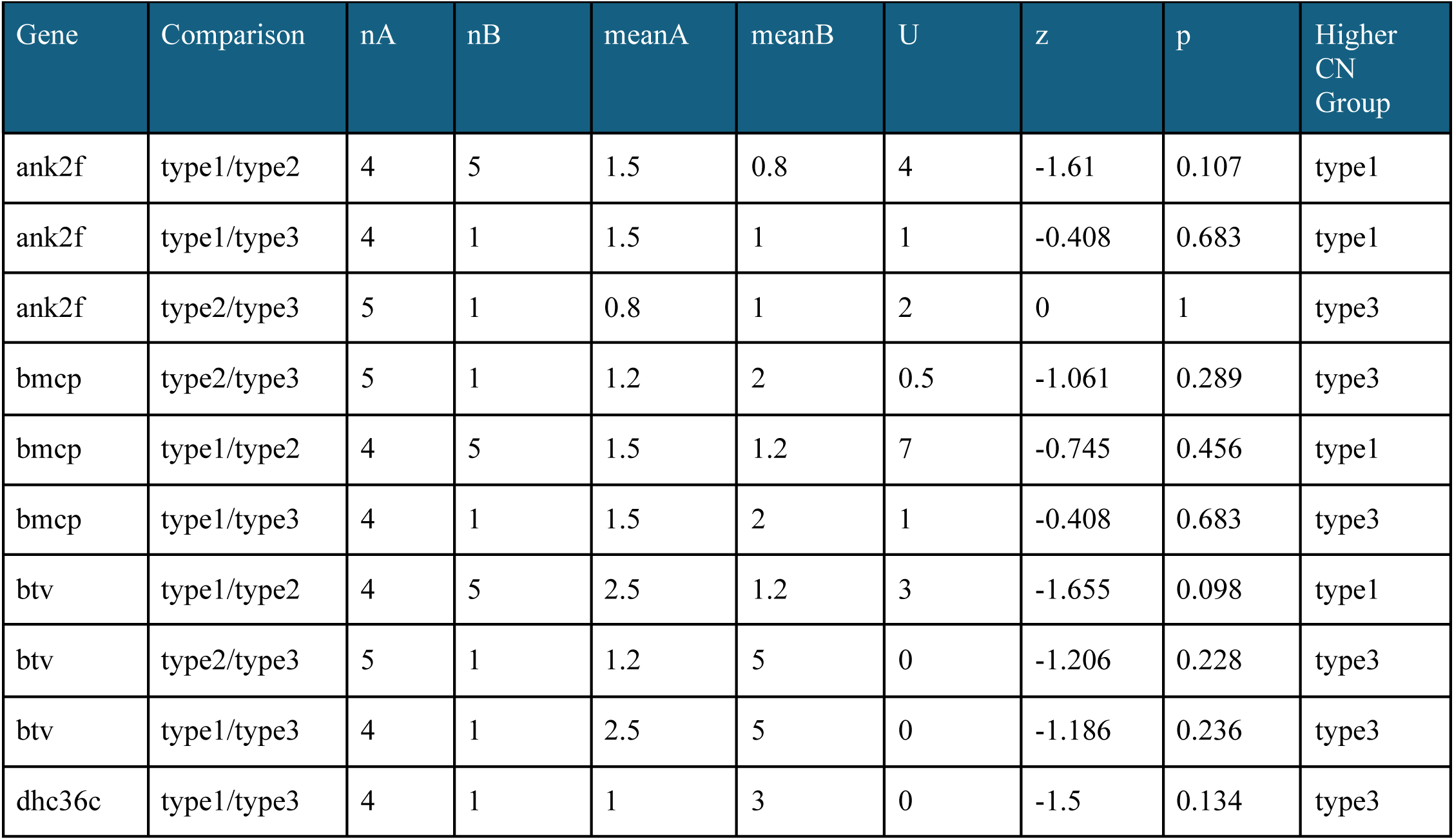
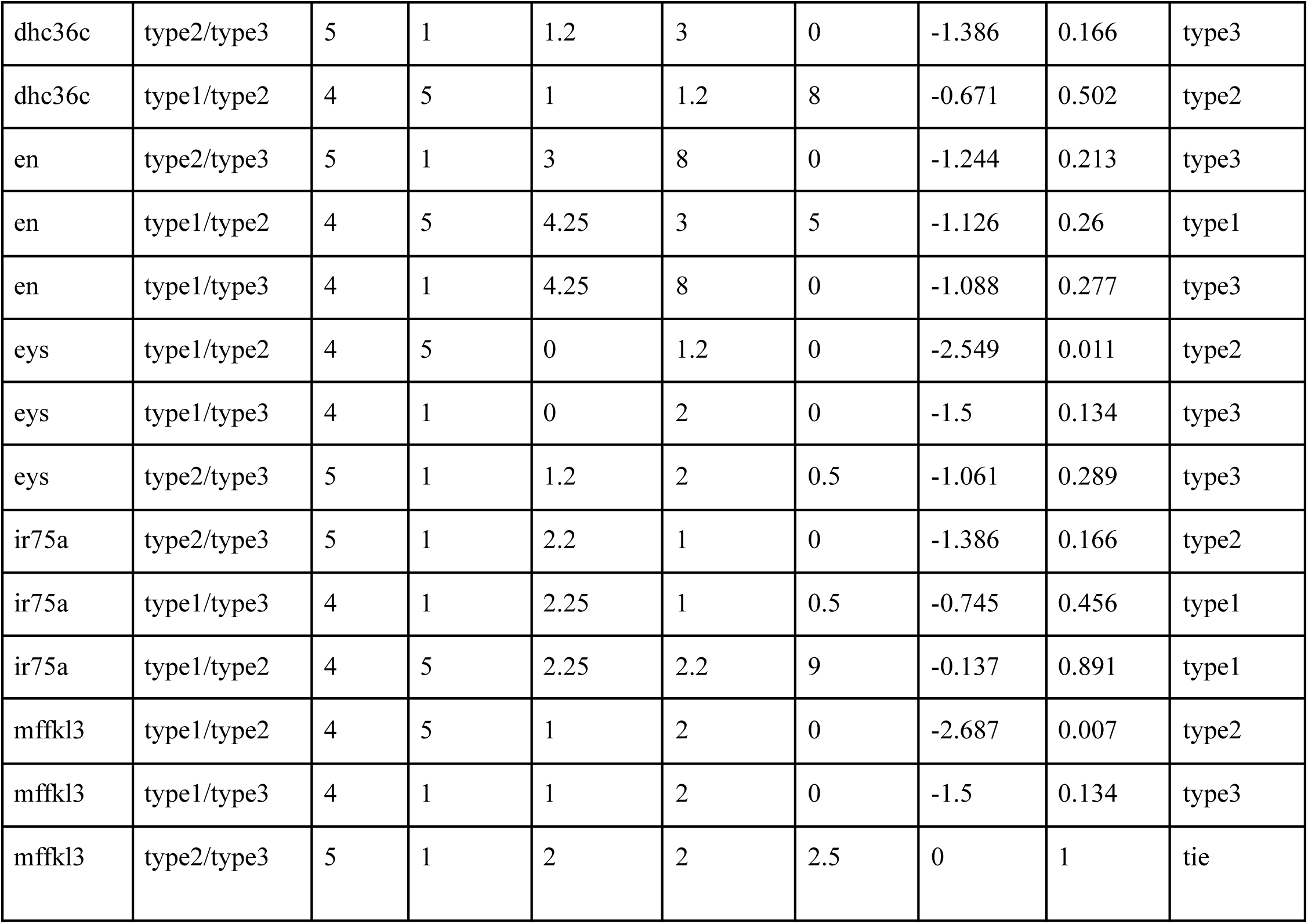
Pairwise Mann-Whitney U test results for comparisons between specific pairs of hearing organ morphologies. For each gene and comparison, nA and nB indicate the size, and meanA and meanB indicate mean copy number of each group. The U statistic, standardized z-score, and two-sided p-value are shown for each group.

| Gene | Comparison | nA | nB | meanA | meanB | U | z | p | Higher CN Group |
| --- | --- | --- | --- | --- | --- | --- | --- | --- | --- |
| ank2f | type1/type2 | 4 | 5 | 1.5 | 0.8 | 4 | -1.61 | 0.107 | type1 |
| ank2f | type1/type3 | 4 | 1 | 1.5 | 1 | 1 | -0.408 | 0.683 | type1 |
| ank2f | type2/type3 | 5 | 1 | 0.8 | 1 | 2 | 0 | 1 | type3 |
| bmcp | type2/type3 | 5 | 1 | 1.2 | 2 | 0.5 | -1.061 | 0.289 | type3 |
| bmcp | type1/type2 | 4 | 5 | 1.5 | 1.2 | 7 | -0.745 | 0.456 | type1 |
| bmcp | type1/type3 | 4 | 1 | 1.5 | 2 | 1 | -0.408 | 0.683 | type3 |
| btv | type1/type2 | 4 | 5 | 2.5 | 1.2 | 3 | -1.655 | 0.098 | type1 |
| btv | type2/type3 | 5 | 1 | 1.2 | 5 | 0 | -1.206 | 0.228 | type3 |
| btv | type1/type3 | 4 | 1 | 2.5 | 5 | 0 | -1.186 | 0.236 | type3 |
| dhc36c | type1/type3 | 4 | 1 | 1 | 3 | 0 | -1.5 | 0.134 | type3 |
| dhc36c | type2/type3 | 5 | 1 | 1.2 | 3 | 0 | -1.386 | 0.166 | type3 |
| dhc36c | type1/type2 | 4 | 5 | 1 | 1.2 | 8 | -0.671 | 0.502 | type2 |
| en | type2/type3 | 5 | 1 | 3 | 8 | 0 | -1.244 | 0.213 | type3 |
| en | type1/type2 | 4 | 5 | 4.25 | 3 | 5 | -1.126 | 0.26 | type1 |
| en | type1/type3 | 4 | 1 | 4.25 | 8 | 0 | -1.088 | 0.277 | type3 |
| eys | type1/type2 | 4 | 5 | 0 | 1.2 | 0 | -2.549 | 0.011 | type2 |
| eys | type1/type3 | 4 | 1 | 0 | 2 | 0 | -1.5 | 0.134 | type3 |
| eys | type2/type3 | 5 | 1 | 1.2 | 2 | 0.5 | -1.061 | 0.289 | type3 |
| ir75a | type2/type3 | 5 | 1 | 2.2 | 1 | 0 | -1.386 | 0.166 | type2 |
| ir75a | type1/type3 | 4 | 1 | 2.25 | 1 | 0.5 | -0.745 | 0.456 | type1 |
| ir75a | type1/type2 | 4 | 5 | 2.25 | 2.2 | 9 | -0.137 | 0.891 | type1 |
| mffkl3 | type1/type2 | 4 | 5 | 1 | 2 | 0 | -2.687 | 0.007 | type2 |
| mffkl3 | type1/type3 | 4 | 1 | 1 | 2 | 0 | -1.5 | 0.134 | type3 |
| mffkl3 | type2/type3 | 5 | 1 | 2 | 2 | 2.5 | 0 | 1 | tie |

**Supplementary Table 7.** Summary of whether a selection test had a significant signal (1) or insignificant signal under three selection tests: aBSREL, BUSTED, and Contrast-FEL.

| Gene Name | Condition Met aBSREL | Condition Met BUSTED | Condition Met Contrast-FEL |
| --- | --- | --- | --- |
| ForkheadDomain | 0 | 0 | 1 |
| DyneinIntermedChain | 1 | 1 | 1 |
| Cut | 0 | 0 | 0 |
| TransReceptorPotIsoB | 0 | 0 | 1 |
| Rhodopsin5 | 0 | 0 | 0 |
| Crinkled | 0 | 0 | 0 |
| Piezo | 0 | 0 | 1 |
| NoMechanoRPotA | 1 | 0 | 0 |
| Inactive | 1 | 0 | 0 |
| Beethoven | 1 | 1 | 1 |
| ReducedMechanoRPotA | 0 | 0 | 1 |
| Arrestin2 | 0 | 0 | 0 |
| Bmcp | 0 | 0 | 0 |
| ScavengerReceptor | 0 | 0 | 1 |
| TouchInsensitiveLarvaB | 0 | 0 | 0 |
| NaPumpAlphaSubunit | 1 | 0 | 0 |
| Atonal | 0 | 0 | 0 |
| Rhodopsin6 | 0 | 0 | 0 |
| CG9492 | 0 | 0 | 0 |
| Waterwitch | 1 | 0 | 0 |
| NoMechanoRPotB | 1 | 1 | 1 |
| TransReceptorPotIsoA | 0 | 0 | 1 |
| Spalt-related | 1 | 1 | 0 |
| Forked | 0 | 0 | 0 |
| Seizure | 0 | 0 | 0 |
| Nanchung | 0 | 0 | 0 |
| CG4329 | 0 | 0 | 0 |
| KinesinAssocProtein | 1 | 0 | 0 |
| Calmodulin | 1 | 0 | 0 |
| MaleFertilityFactor | 0 | 0 | 0 |
| KingTubby | 0 | 0 | 0 |
| Diaphanous | 0 | 1 | 0 |
| TetraspanMembraneHairStereocilia | 0 | 0 | 1 |
| Nervana2 | 0 | 1 | 0 |
| EyesShut | 0 | 0 | 1 |
| TechnicalKnockout | 1 | 0 | 0 |
| Slowterm | 0 | 0 | 0 |
| Brown | 0 | 0 | 0 |
| Rfx | 0 | 0 | 1 |
| DyneinHeavyChain93<br>AB | 0 | 0 | 0 |
| IonotropicReceptor75a | 0 | 1 | 0 |
| CG14636 | 0 | 0 | 0 |
| KinesinLikeProtein | 1 | 1 | 0 |
| Ankyrin2IsoformX | 1 | 0 | 1 |
| RetinalDegenerationA | 1 | 1 | 0 |
| Inactivation | 0 | 0 | 1 |
| CG9313 | 0 | 1 | 0 |
| Ankyrin2IsoformF | 1 | 0 | 1 |
| DyneinHeavyChain36<br>C | 0 | 0 | 0 |
| NoMechanoRPotC | 0 | 0 | 0 |
| Glass | 0 | 0 | 1 |
| Engrailed | 1 | 0 | 1 |
| Sosie | 0 | 0 | 1 |
| ZincFinger | 0 | 0 | 0 |
| DoublecortinDomainP<br>rotein | 0 | 0 | 0 |
| CG8086 | 0 | 0 | 0 |

**Supplementary Table 8.**
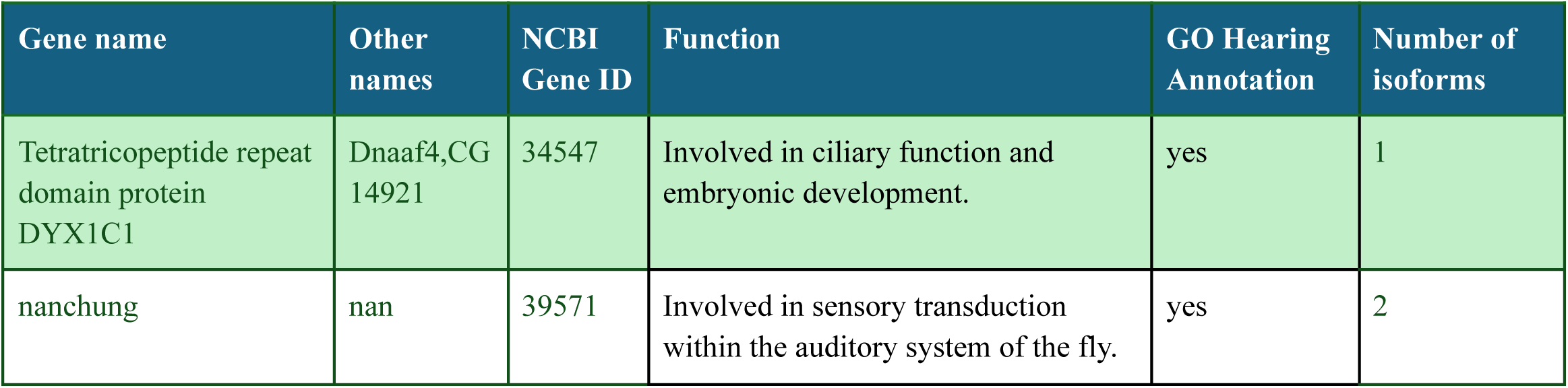
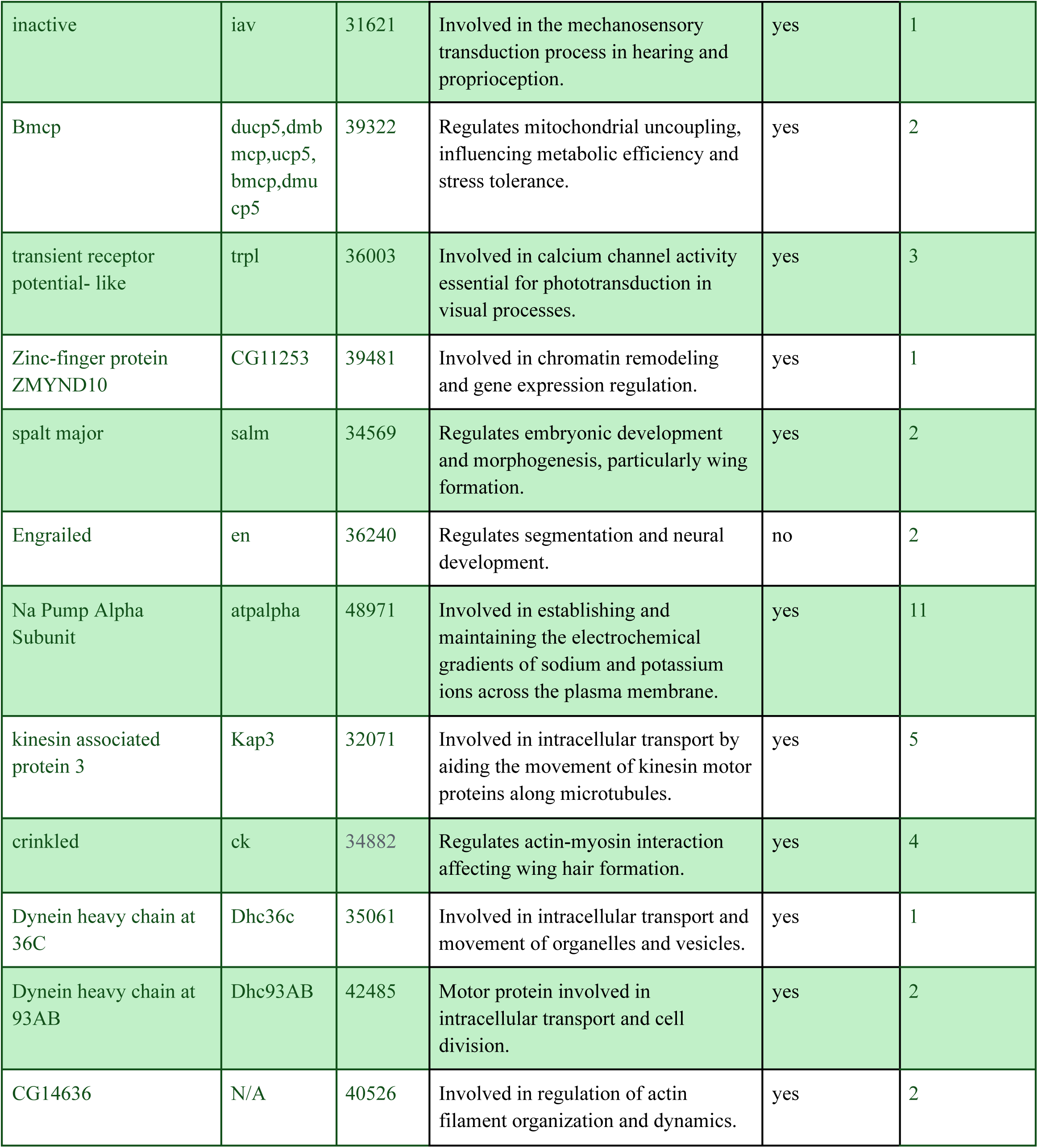
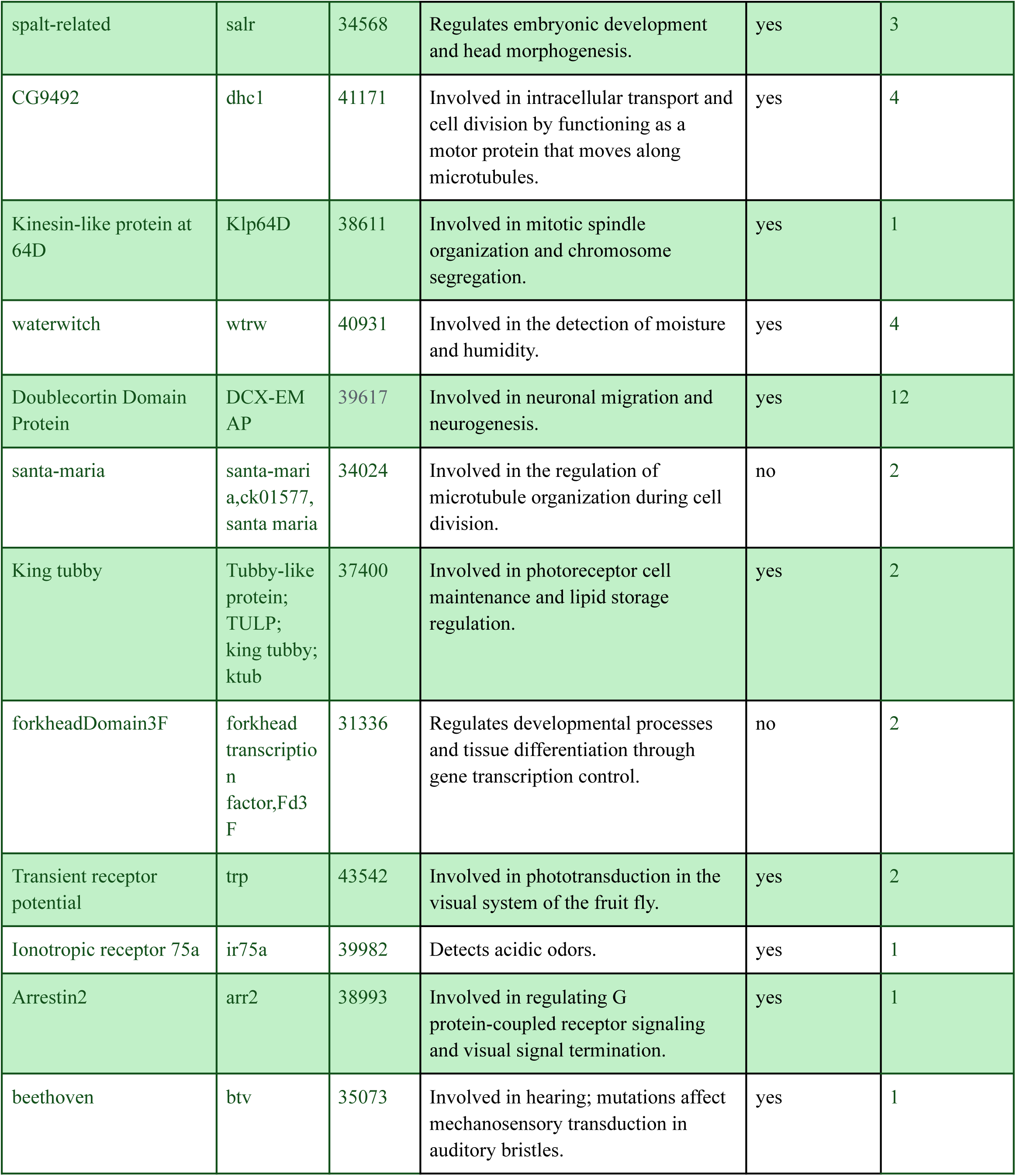
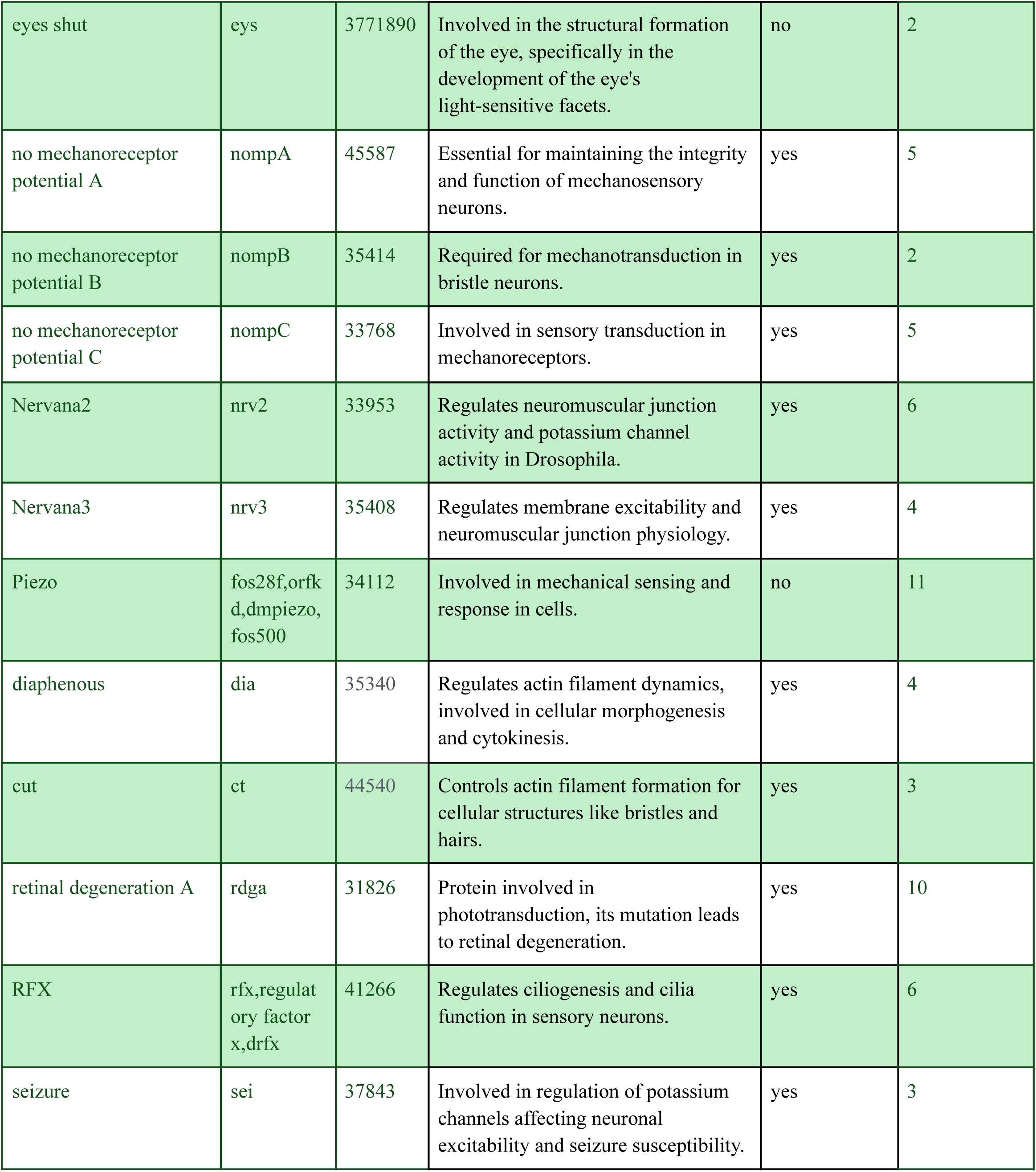
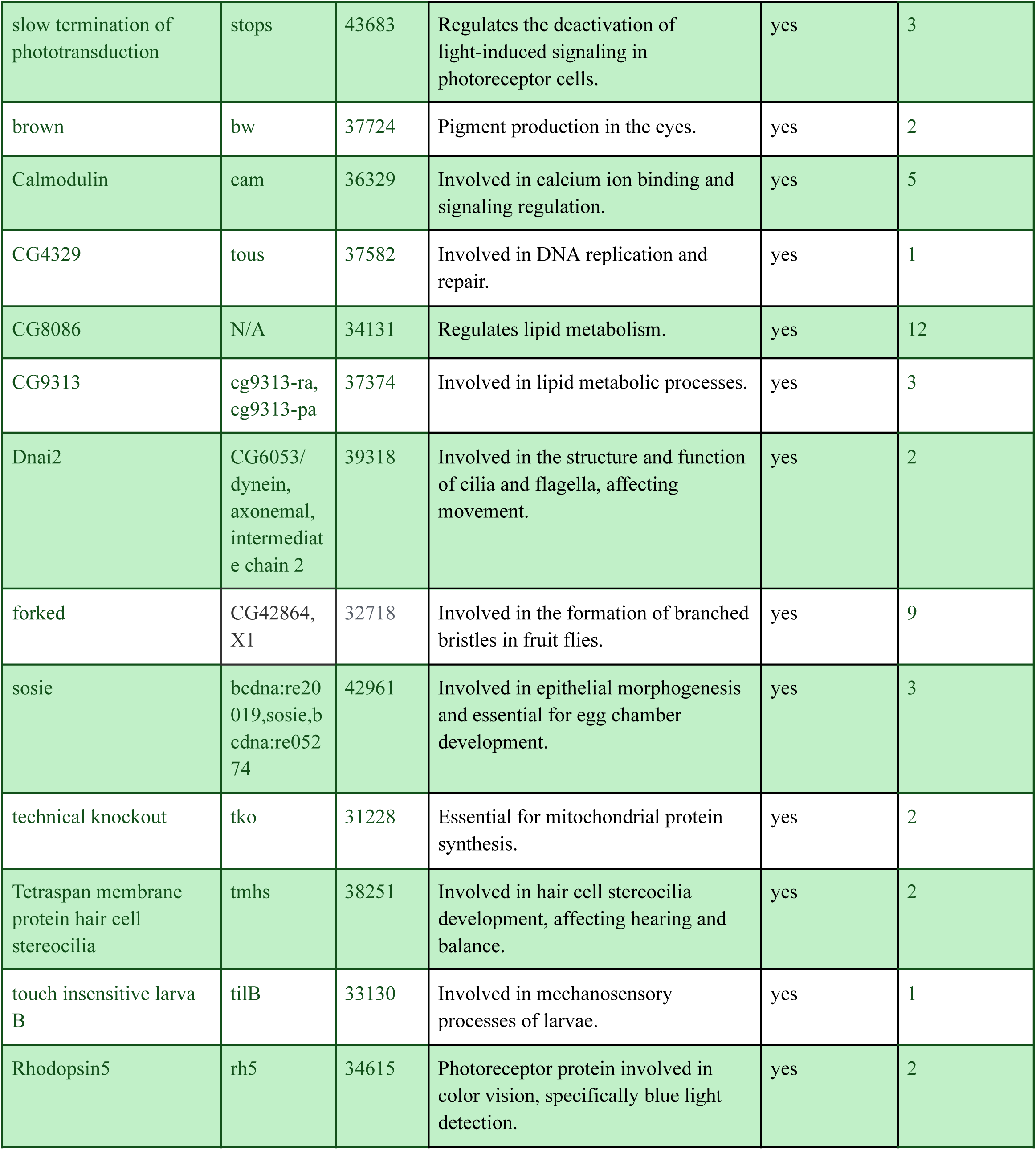
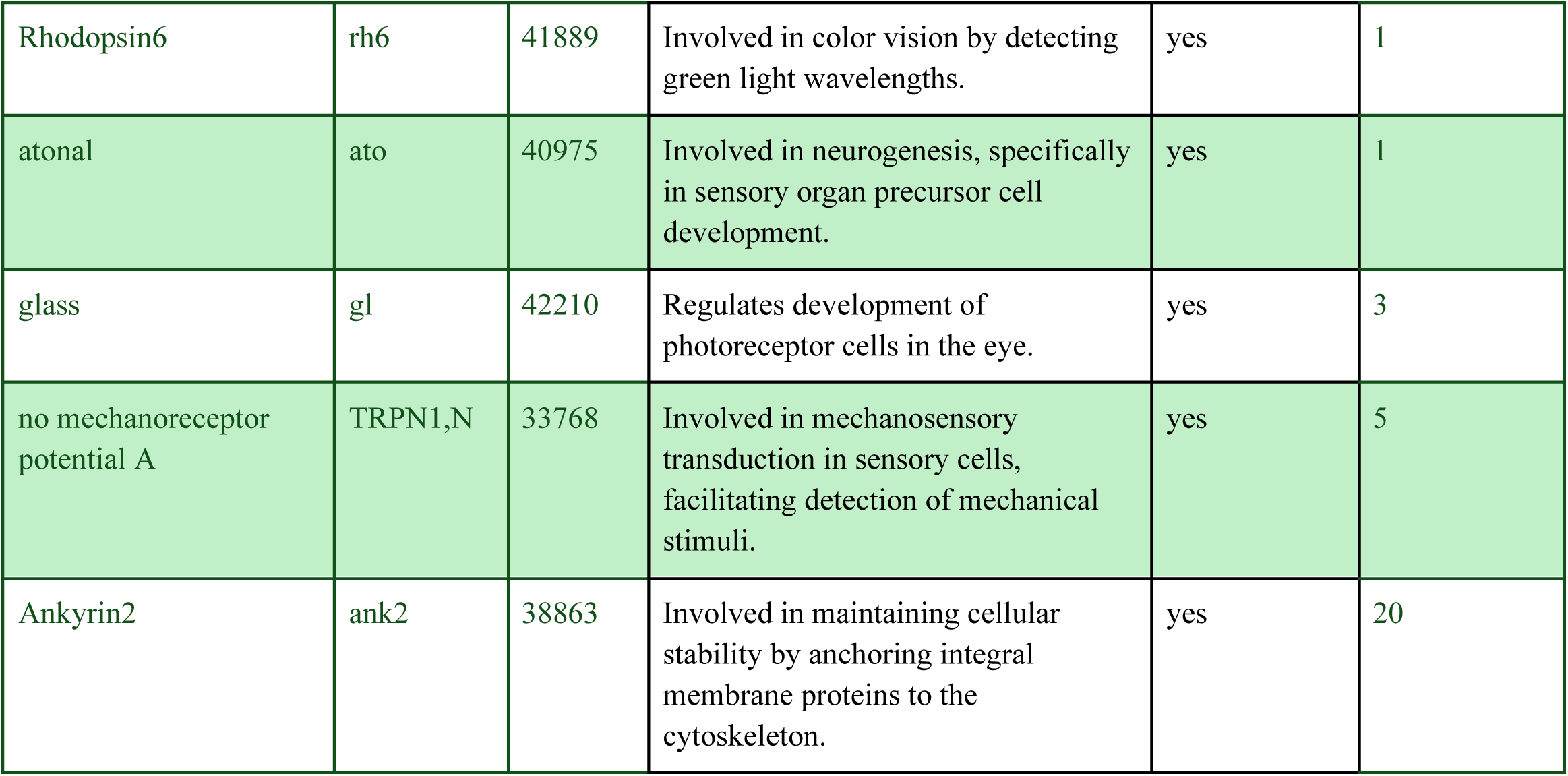
56 genes with frequent alternative names, NCBI Gene ID, function, number of isoforms, and indicate of GO annotation related to hearing.

**Supplementary Table 9.**
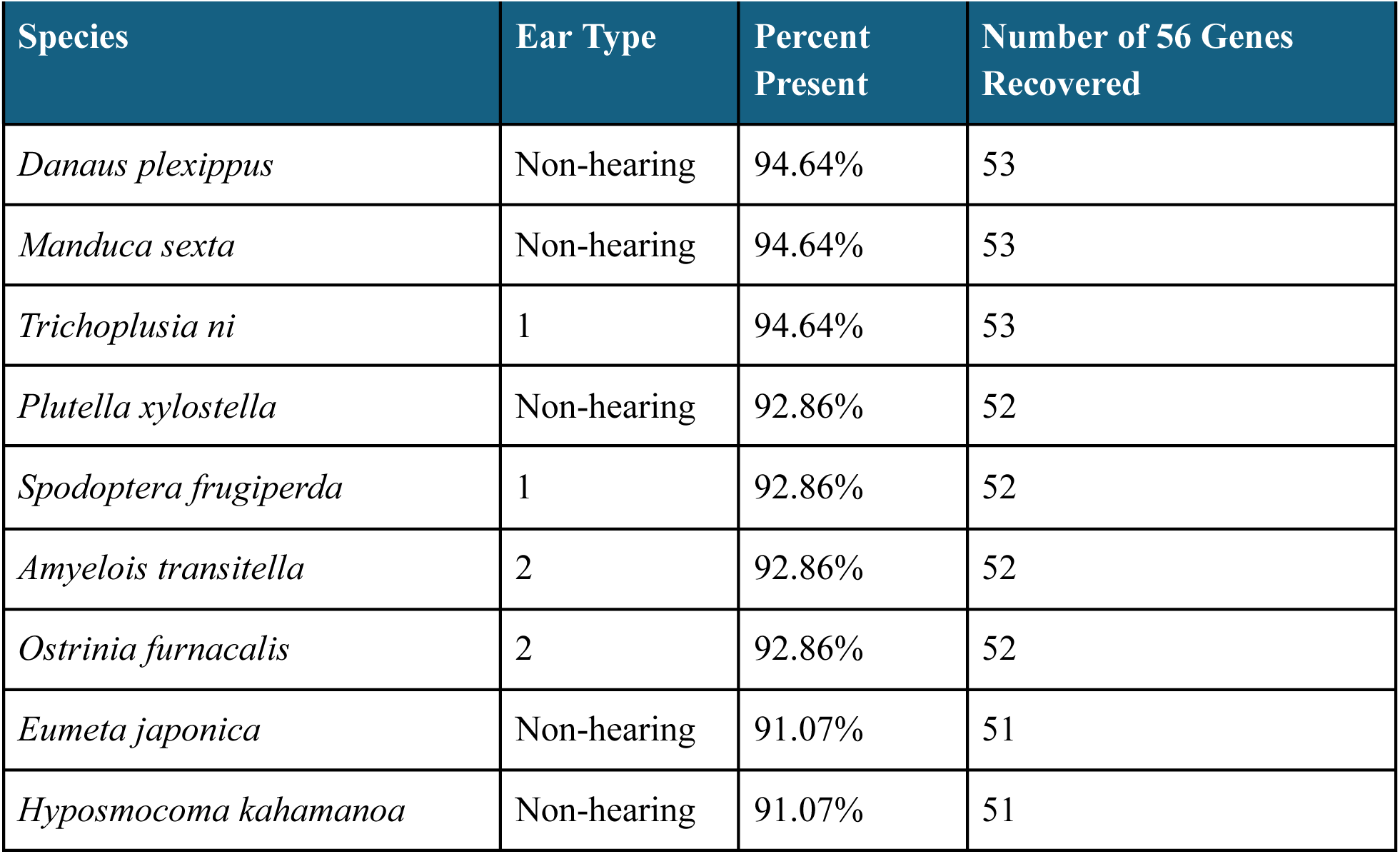
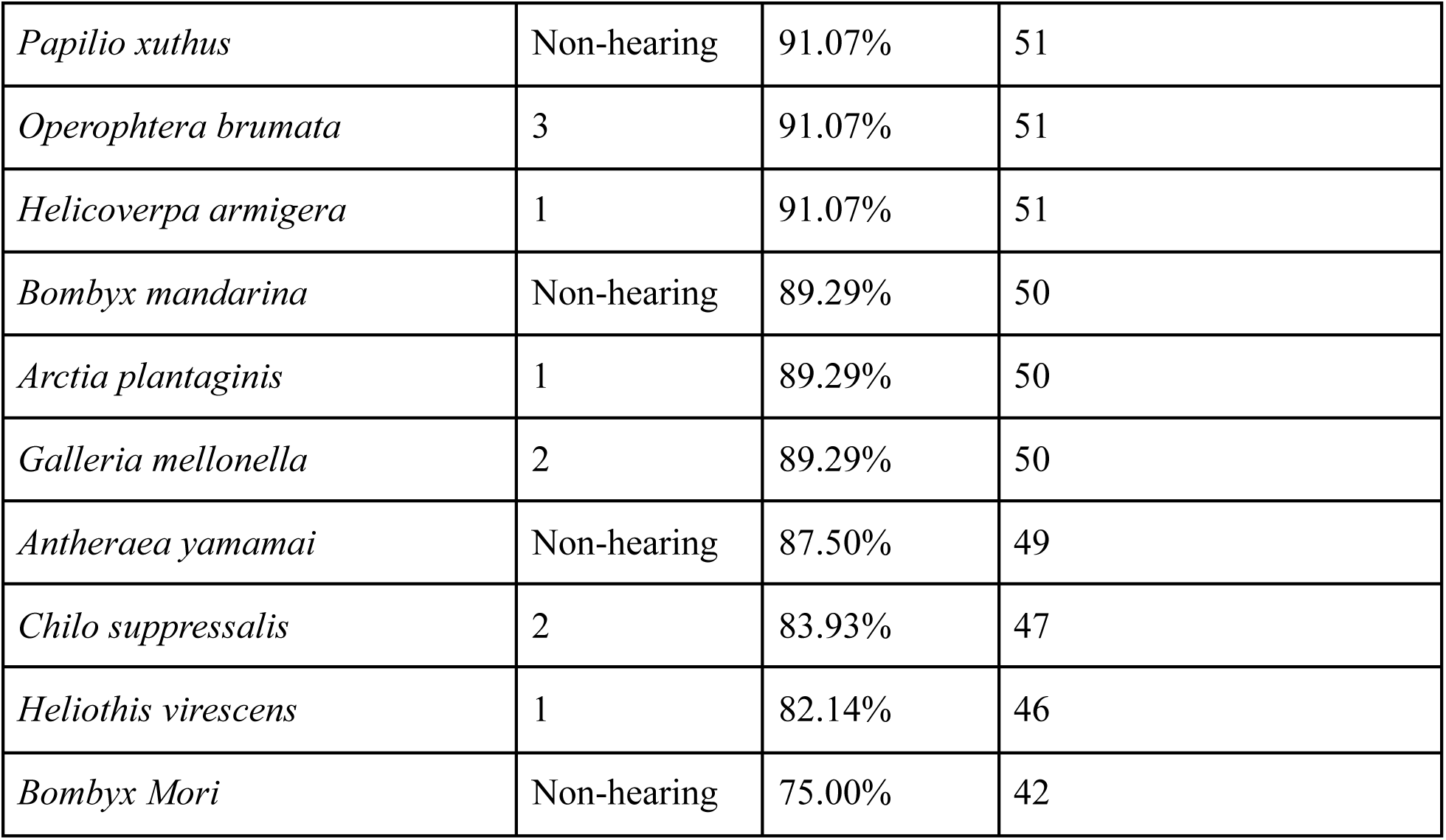
Number and percentage of 56 genes found to have an ortholog in a given species.

## Supplementary Methods

### Supplementary Method 1. Ortholog Filtering and Representative Sequence Selection

After the initial clustering of sequences with OrthoFinder, we filtered sets of candidate orthologs for each gene such that a single sequence was selected for each species for later analysis. An all-vs-all BLAST search was performed within the file and the sequence most representative of the set (least average divergence from other sequences) was selected as assessed by query coverage and percent identity. Sequences were then grouped by species and compared via BLAST against the chosen representative sequence for the highest overall number of matching bases. These selected sequences were used for later alignment and selection tests.

### Supplementary Method 2. Measurement of ω_hearing_ > ω_non-hearing_ and ω_hearing_ < ω_non-hearing_

Contrast-FEL hits were filtered and organized such that only hits with a pOverall statistic <0.05 were considered, and only on genes with >5 codon sites affected. These were filtered to retain only those with a ratio of enriched (ω_hearing_ or ω_non-hearing) over total significant sites, of >0.60. The resulting output is shown in Supplementary Tables 1-2.

